# Mice exhibit stochastic and efficient action switching during probabilistic decision making

**DOI:** 10.1101/2021.05.13.444094

**Authors:** Celia C. Beron, Shay Q. Neufeld, Scott W. Linderman, Bernardo L. Sabatini

**Affiliations:** Howard Hughes Medical Institute, Department of Neurobiology, Harvard Medical School, Boston, MA; Department of Statistics and Wu Tsai Neurosciences Institute, Stanford University, Stanford, CA

## Abstract

In probabilistic and nonstationary environments, individuals must use internal and external cues to flexibly make decisions that lead to desirable outcomes. To gain insight into the process by which animals choose between actions, we trained mice in a task with time-varying reward probabilities. In our implementation of such a “two-armed bandit” task, thirsty mice use information about recent action and action-outcome histories to choose between two ports that deliver water probabilistically. Here, we comprehensively modeled choice behavior in this task, including the trial-to-trial changes in port selection – i.e. action switching behavior. We find that mouse behavior is, at times, deterministic and, at others, apparently stochastic. The behavior deviates from that of a theoretically optimal agent performing Bayesian inference in a Hidden Markov Model (HMM). We formulate a set of models based on logistic regression, reinforcement learning, and ‘sticky’ Bayesian inference that we demonstrate are mathematically equivalent and that accurately describe mouse behavior. The switching behavior of mice in the task is captured in each model by a stochastic action policy, a history-dependent representation of action value, and a tendency to repeat actions despite incoming evidence. The models parsimoniously capture behavior across different environmental conditionals by varying the ‘stickiness’ parameter, and, like the mice, they achieve nearly maximal reward rates. These results indicate that mouse behavior reaches near-maximal performance with reduced action switching and can be described by a set of equivalent models with a small number of relatively fixed parameters.

**Significance:** To obtain rewards in changing and uncertain environments, animals must adapt their behavior. We found that mouse choice and trial-to-trial switching behavior in a dynamic and probabilistic two-choice task could be modeled by equivalent theoretical, algorithmic, and descriptive models. These models capture components of evidence accumulation, choice history bias, and stochasticity in mouse behavior. Furthermore, they reveal that mice adapt their behavior in different environmental contexts by modulating their level of ‘stickiness’ to their previous choice. Despite deviating from the behavior of a theoretically ideal observer, the empirical models achieve comparable levels of near-maximal reward. These results make predictions to guide interrogation of the neural mechanisms underlying flexible decision-making strategies.

## Introduction

Animals select appropriate actions to achieve their goals. Furthermore, animals adapt their decision-making process as the environment changes. During foraging, for example, animals make decisions about when and where to search for food to safely acquire sufficient nutrients. This requires balancing the trade-off between exploiting known sources of food versus continuing to explore unknown, potentially more profitable options. In a dynamic environment, continued exploration and adaptation are required to detect and react to changing conditions, such as the depletion or appearance of a food source, that may influence what decision is optimal at a given time. Inherent in this process is the ability to accumulate evidence about the value of various actions from previous experience. Many neuropsychiatric diseases are associated with perturbations of evidence-dependent action selection (i.e., cognitive and/or behavioral flexibility), making individuals with the disease resistant to updating action plans despite changes in environmental contingencies (reviewed in [1–4]).

The dynamic multi-armed bandit task is an experimental paradigm used to investigate analogs of these decision-making behaviors in a laboratory setting [5–13], including in normotypic humans and those with disease [14–16]. In this task, the experimental subject chooses between a small number of actions, each of which offers a nonstationary probability of reward. The dynamic reward contingencies require the players to flexibly modulate their actions in response to evidence accumulated over multiple trials. Therefore, switching between behaviors is a key component of performing this task. However, analysis of behavior in this task is often reduced to examining the agent’s selection of the higher rewarding port or to the detection of a state transition.

Several classes of models have been used to model behavior in two- or multi-armed bandit tasks, which make different assumptions about the underlying decision-making process and focus on different aspects of the behavioral output. For example, theory-guided ideal observer models assume that agents learn the dynamics of reward contingencies and use Bayesian inference to identify the optimal action on each trial [16–18]. Model-free reinforcement learning strategies [19], like the Rescorla-Wagner model [20], are more algorithmic in nature. Rather than assuming knowledge of reward contingency dynamics, these models maintain a running estimate of the value of different actions [9, 10, 14]. Similarly, drift-diffusion models explicitly model evidence accumulation as an inertial process in order to explain hysteresis in action selection [21–23]. Finally, descriptive models make few assumptions about the information integration process but simply predict future behavior given past actions and outcomes using, for example, a logistic regression [6, 13]. These efforts have provided insight into how simple algorithms can reduce a series of actions and outcomes to features that might be represented in the brain, facilitating the identification of neural correlates of action value and belief state representations [6, 10–12, 24]. Furthermore, by enabling the differentiation of trials in which behavior deviates from the action with the highest expected value, such models have revealed neural activity related to exploration [8, 25]. However, in evaluating these various models, trials in which the animal switches between actions are typically not explicitly considered, and, because these are a small minority of trials, failure to model them correctly has little impact on overall model accuracy across all trials.

Here, we develop a statistical analysis of the relatively infrequent subset of trials in which the agent switches between actions, enabling examination of the features that contribute to the flexible and exploratory components of behavior. We use these models to study mouse behavior in a two-armed bandit task and gain insight into the strategy that animals use to select actions to achieve reward. We find that trial-to-trial action switching is a stochastic component of the behavior and sets theoretical limits on the performance of behavioral models in predicting action choice. Although the optimal agent in this task would perform inference in a Hidden Markov Model (HMM), mouse behavior is not consistent with that of such an agent. Instead, it is better-described by a simple logistic regression using a stochastic action-selection policy. Leveraging the simple form of the logistic regression weights, we reformulate this model as one that recursively updates a single state estimate. This recursively formulated logistic regression model not only captures mouse choice and switching behavior but generalizes to new environmental parameters through a parsimonious solution that minimally reduces expected rewards. We further show that this model closely resembles a Q-learning algorithm from reinforcement learning, and under further assumptions they can be shown to be equivalent. Finally, we relate these models to a ‘sticky’ agent performing HMM inference. Altogether, our results connect descriptive, algorithmic, and theoretically motivated model formulations to offer multiple views on animal behavior and make predictions about its underlying neural mechanisms.

## Results

### Task structure and performance

To study probabilistic decision making, we trained mice in a Markovian two-armed bandit task. During each behavior session, the mouse moved freely in a chamber containing three ports into which it could place its snout (i.e. nose poke) to engage with the task (Fig. 1A). One of the side ports delivered reward with probability *p* ∈ [0.5, 1] (the “high” port) and the other with probability 1− *p* (the “low” port). We trained each mouse with three sets of task conditions, in which the high-low reward probabilities (in percent) were assigned as 90-10, 80-20, or 70-30 for a given session but changed day-to-day. The state of the reward probabilities was assigned on a trial-by-trial basis following a Markovian process, such that after completion of each trial high and low ports remained the same with probability *q* = 0.98 and switched with probability 1− *q* = 0.02. This stochastic process produced blocks of consecutive trials during which the high reward probability was assigned to the right or left port (Fig. 1B), with a mean block length of 50 trials.

**Figure 1:**
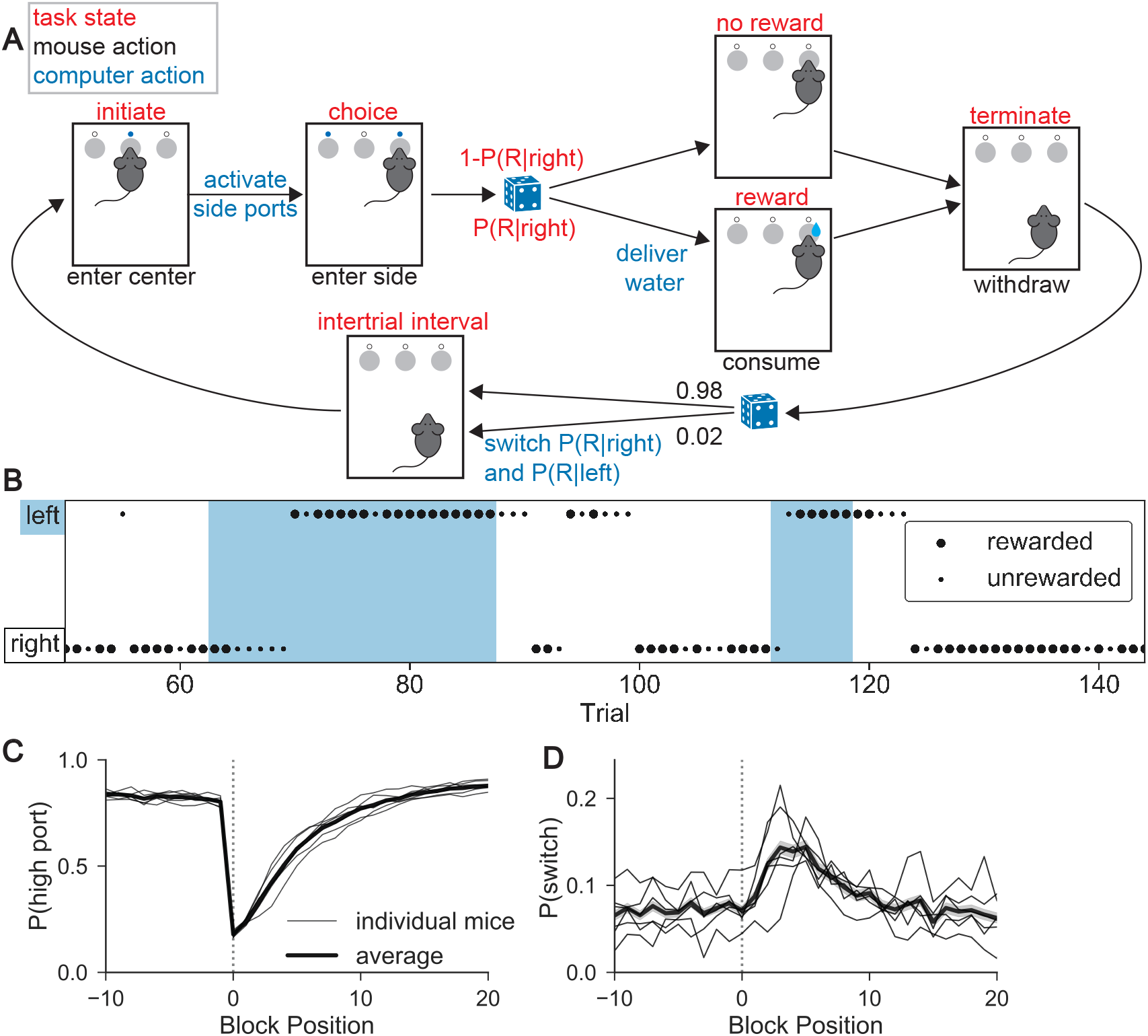
Mouse behavior in a two-armed bandit task. (A) Task structure: A mouse initiates a trial by putting its snout (i.e. “poking”) into the center port. It then selects one of the two side ports in order to enter the “choice” state. In this illustration, the mouse chose the right port. Depending on the choice and preassigned port reward probabilities, reward is or is not delivered. The mouse “terminates” the trial by withdrawing from the side port, which initiates the “inter-trial interval” state. During this 1 s period, the computer assigns reward probabilities for the subsequent trial using a Markov process. (B) Example mouse behavior across part of a session. Blue and white shading indicates the location of the high reward probability port as left and right, respectively. Dot position and size indicate the port chosen by the mouse and the outcome of the trial, respectively (large = rewarded). (C) *P*_highchoice_ for *p* = 0.8 as a function of trial number surrounding the trial at which the reward probabilities reverse (block position=0). Each thin line shows the behavior of an individual mouse (N=6) whereas the thicker line and the shading around it show the mean and standard error, respectively, across mice. (D) As in (C) but for *P*_switch_.

Wild-type mice learned to perform this task in all three sets of reward conditions. We focus on the intermediate condition, 80-20, in the main text and figures unless otherwise stated, but corresponding information for the alternative contexts is reported in the supplementary materials. In the 80-20 sessions, mice achieved an average of 514±77 water rewards in a 40 min session (±SD, n=6, Table S1). Overall, right and left port selection was unbiased (51% left, 49% right) and mice performed each trial quickly (center port to center port elapsed time or trial durations of mean ± SD = 2.05 ± 3.14 s and median ± MAD = 1.65 ± 0.79 s). The mean time between center and choice port was 0.47 s, much faster than the 2 s upper limit imposed by the task structure. Although timing of actions was history dependent (Fig. S1), this information was not used in the analyses and models presented below.

To quantify task performance and characterize the behavioral strategy, we determined the per trial probabilities of (1) selecting the higher rewarding port (*P*_highchoice_), reflecting the ability of the mouse to collect information across trials to form a model of the optimal action, and of (2) switching port selection from one trial to the next (*P*_switch_). Switch trials occurred infrequently: in the 80-20 sessions they made up only 0.07 of all trials. Mice made decisions in a clearly non-random pattern: across mice, *P*_highchoice_ was 0.83 (range: 0.81-0.84, Table S2). Furthermore, the strategy employed by the mice deviated from a simple “win-repeat, lose-switch” strategy as *P*_switch_ was 0.02 following rewarded choices and 0.18 following unrewarded choices (as opposed to the 0.0 and 1.0 rates predicted by win-repeat, lose-switch, Table S1).

Mice were sensitive to the nonstationary reward probabilities: they generally chose the higher rewarding port but adjusted their behavior in response to reward probability reversals at block transitions (Fig.1C). The mice required multiple trials to stably select the new high reward probability port after a block transition (*τ* = 4.28 ± 0.19 s.e.m. trials) (Fig. 1C, Table S2). Furthermore, although across all trials *P*_switch_ was low, it increased after the block transition (Fig. 1D), paralleling the recovery of *P*_highchoice_. The dynamics of *P*_highchoice_ and *P*_switch_ following the block transitions indicate that mice, as expected, modulate their behavior in response to the outcomes of choices and motivates our pursuit of models that capture this behavioral strategy [6, 7, 10, 11]. Mice adapted their behavior across reward contexts, responding more quickly to block transitions in sessions with the more deterministic reward probabilities (90-10) than in those with the more stochastic reward probabilities (70-30) (Fig. S2, Table S2).

### History dependence of behavior

To examine the contribution of trial history to mouse choice, we computed the conditional probability that the mouse switched ports given each unique combination of choice-reward sequences in the preceding trials (P(switch | sequence)), akin to *n*-gram models used in natural language processing [26, 27]. This can be thought of as a nonparametric policy in which the combination of previous choices and rewards (implicitly across varying latent states) guides future choice (Fig. 2A). We used a code to represent the conditioned history sequences, which fully specifies port choice and action outcome over a chosen history length (3 in the given example) leading up to each trial (Fig. 2B, Methods). For a history length of 3 trials, switching behavior has left-right symmetry. For example, the probability of a left choice following three rewarded left choices is approximately the same as the probability of a right choice following three rewarded right choices. This allowed us to represent choice direction in relative terms (Fig. 2C).

**Figure 2:**
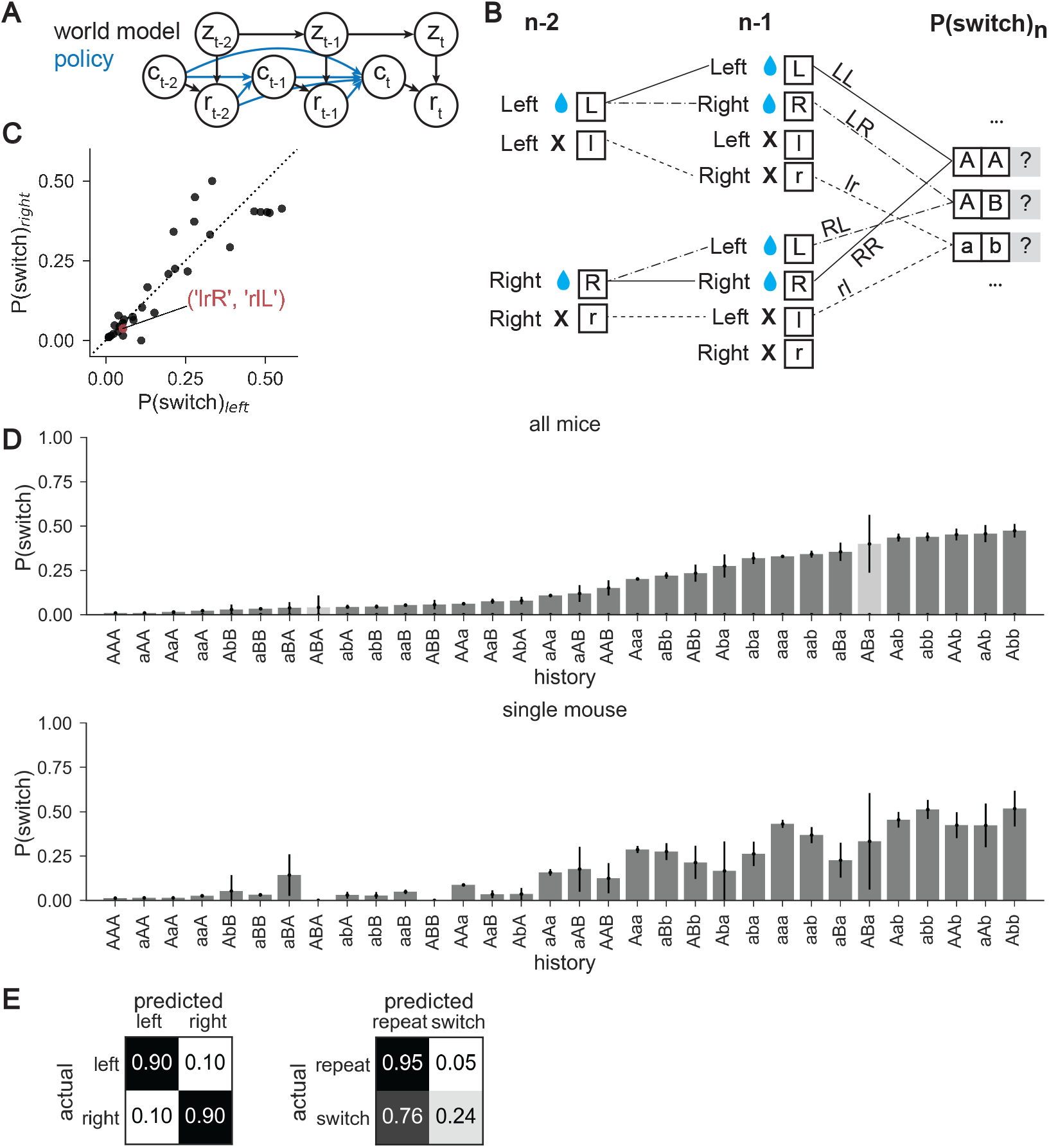
Switching behavior is probabilistic and history dependent. (A) Schematic of world model (black lines) for the two-armed bandit task: rewards (*r*) depend on mouse choice (*c*) and the underlying state (*z*) for each trial (*t*). World state evolves according to a Markov process. A nonparametric policy (blue) shows previous choices and rewards contributing to future choice. (B) The action-outcome combination for each trial is fully specified by one of four symbols: “L” or “R” for left or right rewarded trial, “l” or “r” for left or right unrewarded trial, respectively. These can form “words” that represent action-outcome combinations across sequences of trials. Each sequence starting with right port selection has a mirror sequence starting with a left port selection (e.g. r-L and l-R, in panel A) and can be combined by defining the initial direction in the sequence as “A/a” and those in the other direction as “B/b.” The probability of switching ports on the next trial is calculated, conditioned on each trial sequence for history length *n*. (C) The conditional switch probabilities after R/L mirror pairs of history length 3 are plotted for histories starting on the left vs. right port. The clustering of points around the unity line confirms the symmetry of mouse switching (correlation coefficient = 0.91). One such pair (l-r-R and r-l-L) is highlighted, which becomes a single sequence (a-b-B). (D) top: Conditional switch probability across all mice in the 80-20 condition for each action-outcome trial sequence of history length 3, sorted by switch probability. Each bar height indicates the mean switch probability following the corresponding action-outcome history across all trials and mice. The error bars show binomial standard errors. Sequences that occur with s.e.m.>20% are shown in lighter gray. bottom: As above for data collected across all sessions for a single representative mouse. Sequences are presented along the x-axis using the same order as in the top graph. (E) Confusion matrices for the nonparametric policy for right and left port choice (*left*) and repeat and switch (*right*) in the 80-20 condition. On-diagonal values represent the theoretical maximum for sensitivity, or the proportion of predicted positives relative to all positives, under the mouse’s conditional probability distribution. Off-diagonal values represent expected proportion of false negatives, normalized to one across the row with true positives.

### Apparent stochasticity of behavior limits the accuracy of predictive models

To characterize the history dependence of the mouse switching behavior, we examined conditional switch probabilities for all unique action and outcome sequences for history length 3 (Fig. 2D, Fig. S2). This showed that the probability of switching varies as a function of trial history, supported by cross-validated likelihood estimates on data held-out from the ∼115,000 trials collected from the 80-20 sessions (See Methods, Fig. S3), and confirms that mouse behavior depends on action and outcome history. Broad trends can be identified such as the tendency to repeat the previous action after rewarded trials. In addition, although mice exhibit a regime of behavior in which they nearly deterministically repeat the same port choice on subsequent trials (*P*_switch_ ≈ 0), the maximum conditional *P*_switch_ does not approach 1 for any action/outcome sequence, instead reaching a maximum of ∼0.5 (P(switch | “Abb”) =0.47 ± 0.078 s.e.m.). Thus, switches cannot be predicted with certainty for any combination of 3 past actions and outcomes. This apparent stochasticity persists for longer history sequences that are expressed sufficiently often to reliably calculate P(switch | sequence) (Fig. S2-3). Thus, mouse behavior can, in this framework, be qualitatively described as moving from an “exploit” state of repeating recently rewarded actions to an “explore” state of random port choice after recent failures to receive reward.

For a history of length 3, this nonparametric model of mouse behavior is defined by 4^3^/2 = 32 conditional probabilities. A more concise summary is given by the confusion matrices for its average predicted choice probabilities (Fig. 2E). We considered two representations of these choices: the chosen port (left/right) and whether the mouse switched port from the last trial (repeat/switch). These confusion matrices show that left and right port choice are highly predictable actions, each with an average probability of 0.90. In contrast, although the repetition of action selection from one trial to the next is highly predictable given choice and outcome history, with an average probability of 0.94, the apparently stochastic nature of switching events makes them highly unpredictable, such that the probability of predicting that the mouse will switch its port choice from one trial to the next is only 0.23. Nevertheless, this prediction is better than that expected by chance given the 0.07 basal switch rate, providing a target against which we can evaluate model performance.

### Models of mouse behavior

Our goal in the preceding analysis of mouse behavior was to identify quantifiable features that could be used to constrain and test computational models of behavior. Based on this analysis we selected four criteria to evaluate models of mouse behavior:

1. The average log likelihood (LL) of a held-out fraction (30%) of data; i.e., the average log probability the model assigns to a mouse’s choice given its preceding choices and the rewards conferred. As a baseline, we use the log likelihood under the nonparametric model with history length 3 (LL = -0.180, Table S3).
2. The ability of the model to accurately predict port selection and switching events on a trial-by-trial basis, as compared to the expected confusion matrices defined above (Fig. 2E).
3. The ability of the model to capture the conditional action and outcome history dependence of *P*_switch_, including the apparent history-dependent stochasticity of behavior (Fig. 2D).
4. The ability of the model to reproduce the dynamics of *P*_highchoice_ and *P*_switch_ around block transitions (Fig. 1C-D).

These features of behavior were stable within and across sessions (Fig. S4, S5).

In developing models, we separately consider two components underlying the observed behavior: the algorithm and the policy. The algorithm is the process used to generate ‘beliefs’ about the state of the environment (i.e. level of confidence that the higher reward port is left vs. right). The policy relates those computed beliefs to a decision to select a port. The behavioral task evolved according to a discrete Markovian process such that, from the agent’s perspective, the world can be described as governed by a Hidden Markov Model (HMM). Therefore, the theoretically motivated, ideal observer model would use a Bayesian inference algorithm for HMMs to infer which port is most likely to yield reward. We compared this theoretically motivated model to logistic regression—a descriptive model that is frequently used to predict behavior in this context—and to Q-learning—a commonly used reinforcement learning algorithm. For the policy, we hypothesized that stochastic action policies would better reproduce the observed behavioral patterns over their deterministic counterparts, given the apparent stochasticity of mouse conditional switch probabilities.

### Bayesian agents fail to capture mouse behavior

In our task, there are two environmental states corresponding to whether the left or right port is the higher reward probability port. These states are not directly observable by the mouse. Instead, they are relayed to the mouse through the outcomes of its choices. A Bayesian agent computes a posterior distribution (also called the “belief state”) over the environmental state given past choices and rewards by performing inference with a model of the world, here a Hidden Markov Model (HMM). Due to the Markovian nature of the task, the belief state computation can be performed recursively (Fig. 3A, Methods). The agent then incorporates the belief state into a policy, which specifies a distribution over choices on the next trial.

**Figure 3:**
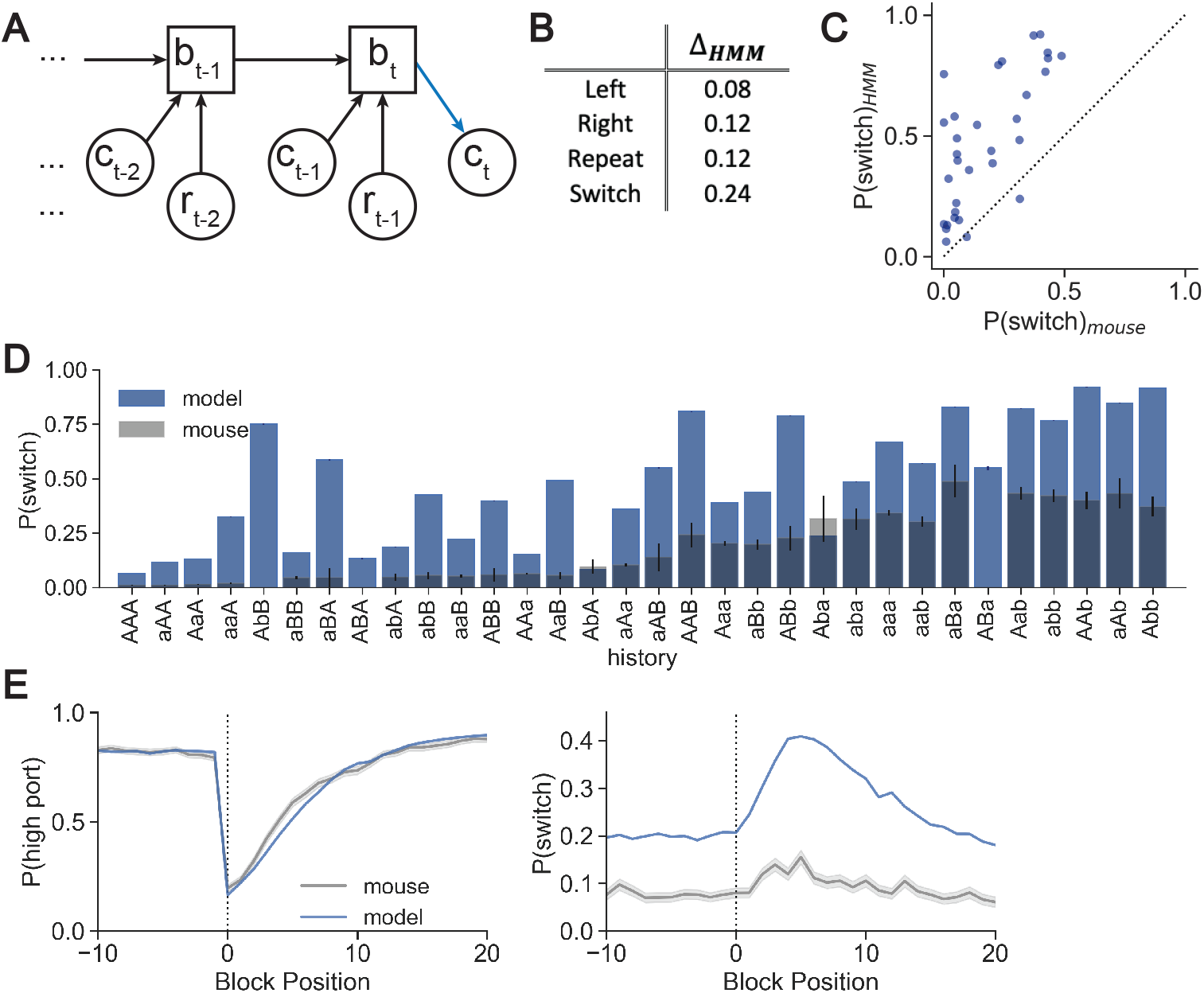
Hidden Markov model overestimates mouse switching probability. (A) The Hidden Markov model (HMM) recursively updates belief state (*b*_*t*_) by incorporating evidence from choice (*c*_*t* − 1_) and reward (*r*_*t* − 1_) of the recent trial. The next choice (*c*_*t*_) depends on the model posterior and the policy (blue). (B) Absolute values of the differences between the HMM confusion matrices and nonparametric confusion matrix (Fig. 2E) for each action type. (C) Conditional switch probabilities generated from the HMM plotted against those observed from mice (SSE = 4.102). (D) Conditional switch probabilities as predicted by the HMM (blue, ‘model’) overlaid on the observed mouse behavior (gray) for all history sequences of length 3. Sequences on the x-axis are sorted by increasing *P*(*switch*)_*mouse*_ for the full dataset (Fig. 2D). The bar heights show the mean switch probability across mice for each corresponding sequence history, and the error bars show the binomial standard error for the mouse test data. (E) HMM-generated *P*_highchoice_ (blue, *left*) and *P*_switch_ (*right*) as a function of trial number surrounding state transition (block position 0) as compared to the mouse behavior (gray). Dark lines show the mean across trials at the same block position and the shading shows the standard error.

Let *z*_*t*_ denote the environmental state on trial *t* (left: *z*_*t*_ = 1, right: *z*_*t*_ = 0). Let *c*_*t*_ denote the mouse’s choice (left: *c*_*t*_ = 1, right: *c*_*t*_ = 0), and let *r*_*t*_ be a binary variable indicating whether or not the mouse received a reward. Since the environmental state is binary, we can represent the belief state with a single value, *b*_*t*+1_ = *P*(*z*_*t*+1_ = 1 | *c*_1:*t*_, *r*_1:*t*_). For Bayesian agents, the distribution of the next choice is determined by the policy, which is a function of the belief state, *P*(*c*_*t*+1_ = 1 | *c*_1:*t*_, *r*_1:*t*_) = *π*(*b*_*t*+1_).

We considered multiple policies to convert the belief state into a distribution over choices on the next trial. In this task, the optimal agent would use a greedy policy in which *π*(*b*_*t*+1_) = 1 if *b*_*t*+1_ ≥ 0.5 and *π*(*b*_*t*+1_) = 0 otherwise. Alternatively, the Thompson sampling policy [28] sets *π*(*b*_*t*+1_) = *b*_*t*+1_ so that ports are chosen at a rate proportional to the model’s belief. Lastly, the “softmax policy” interpolates between these two policies by means of a temperature parameter, *T* ([29], see Methods). As *T* goes to zero, the softmax policy recovers the greedy policy, and when *T* = 1 it is equivalent to Thompson sampling.

To test if a Bayesian agent could accurately model the mouse’s behavior, we performed a dense grid search over the HMM parameters and selected the parameters that maximized the log probability of the mouse’s choices. We did this for a range of softmax policy temperatures. For the Thompson sampling policy (*T* = 1), the best-fit HMM parameters accurately capture the temporal structure of the environment (maximized at a transition probability of 0.02) but underestimates the high port reward probability (maximized at a reward probability of 0.65 whereas the true probability was 0.8). In terms of predicting the mouse’s behavior, this model was much worse than the baseline (LL = -0.325, Table S3).

We also examined behavior predicted by the Bayesian agent with the Thompson sampling policy and found that it failed to capture essential features of the mouse behavior, as measured by criteria 2-4 above.

This agent systematically overestimated the probability of switching (Fig. 3B-E). This is reflected by the deviation of the model from the expected confusion matrices of the nonparametric policy, which we compute as the absolute values of the differences between the model’s values and expected values for each action (Fig. 3B Δ*s*, compared to the data in Fig. 2E). Accordingly, the model overestimates the conditional switch probabilities. (Fig. 3C-D). (Note: here we present the analyses of the held-out data not used for training, which is only 30% of the data presented in Figure 2D. We preserve the sorting order from the full dataset, but for this reason the conditional switch probabilities and binomial standard error estimates differ across figures.) Finally, the HMM fails to capture the dynamics of *P*_switch_ around block transitions of reward probabilities (Fig. 3E). On the other hand, *P*_highchoice_ is captured quite well by the HMM, demonstrating the ability of a model to predict port selection from action-outcome history despite using very different trial-by-trial switching dynamics than the animal.

We performed the same procedure at different softmax policy temperatures, but by each of the behavioral metrics outlined above, these models also failed to capture the mouse behavior (Fig. S6, Table S3). This included an HMM using parameters that correspond to the ideal observer, which uses a greedy policy wherein the agent deterministically selects the port that has a higher probability according to the model’s belief. These results show that the mouse behavior is not captured by the optimal agent, nor by a model following the same inference process but with imperfect learning of environmental parameters.

### Logistic regression with a stochastic policy better predicts mouse behavior

Logistic regression has been used previously to predict rodents’ choices in similar tasks [6, 7, 11, 13], but its ability to predict trial-by-trial switches has not been evaluated. We built a logistic regression model for the conditional probability of the mouse’s next choice given its past choices and rewards, *P*(*c*_*t*+1_ = 1 | *c*_1:*t*_, *r*_1:*t*_) = *σ*(*ψ*_*t*+1_), where *σ*(*x*) = (1 + *e*^−*x*^)^−1^ is the logistic function and *ψt*+1 are the log-odds. We modeled the log-odds as,

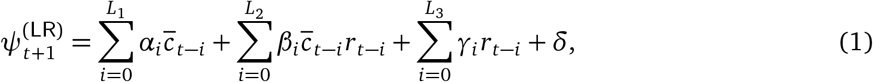

where *α, β*, and *γ* represent the weights on input features for choice 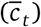, encoding of choice-reward interaction 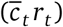, and reward (*r*_*t*_) across trials back to *L*_1_, *L*_2_, and *L*_3_, respectively. The choice is encoded as 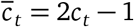, which equals +1 for a left port choice and -1 for a right port choice. We fit the model by maximum likelihood estimation and used cross validation to select the number of past trials to include for each feature. This confirmed that there is minimal left-right port choice bias (i.e., *δ* = 0.04). We also found that rewards alone did not contribute significantly to choice prediction (i.e., *L*_3_ = 0) but that the history of choice-reward encoded trials benefited the model (i.e., *L*_2_ = 5, Fig. 4B). Furthermore, only information about the most recent port choice was necessary (i.e., *L*_1_ = 1). This enabled us to use a reduced form of the model log-odds computation:

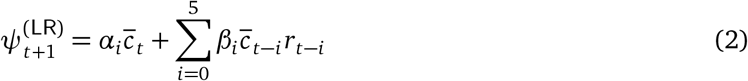

**Figure 4:**
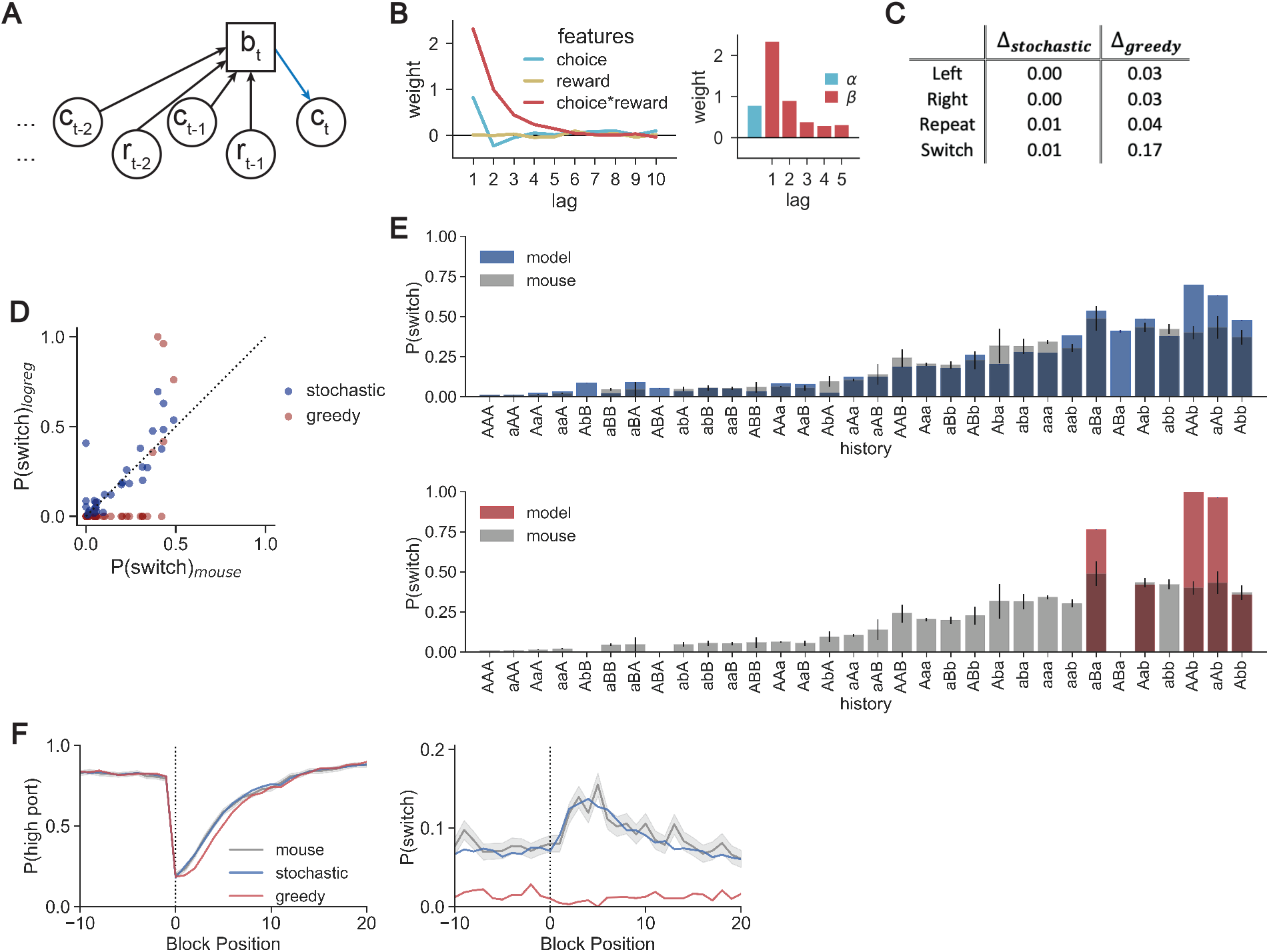
Stochastic logistic regression policy captures mouse behavior comprehensively, whereas greedy logistic regression fails to predict switches. (A) The logistic regression computes the probability of choice (*b*_*t*_) from choice (*c*_*t*− *i*_) and reward (*r*_*t*− *i*_) information across a series of trials. Here we represent the model estimate as *b*_*t*_ for consistency across graphical representations, but note that it in this case it corresponds to the log-odds of choice, *ψ*, in the text. (B) *Left*: Feature weights for a logistic regression predicting the log-odds of mouse port selection for the choices, rewards, and the choice-reward interactions in the previous 10 trials. *right*: Feature weights after cross-validation for hyperparameters and refitting the model. *α* is the weight on the previous choice, and *β* is the set of weights on choice-reward information for the previous 5 trials. (C) Absolute value of the differences between the logistic regression confusion matrices and nonparametric confusion matrix (Fig. 2E) for each action. Δ scores are shown for stochastic logistic regression as well as for greedy logistic regression. (D) Conditional switch probabilities generated by the logistic regression model using a stochastic (blue) or greedy (red) policy plotted against those observed in mice (stochastic-SSE = 0.378, greedy-SSE = 1.548). (E) *Top*: Conditional switch probabilities for the stochastic logistic regression (blue) across sequences of history length 3 overlaid on those from the mouse data (gray). Sequences on the x-axis are sorted according to mouse conditional switch probabilities of the full dataset (Fig. 2D). Error bars show binomial standard errors for the mouse. *bottom*: As above but for a greedy policy (red). (F) *P*_highchoice_ (*left*) and *P*_switch_ (*right*) as a function of trial number surrounding state transition (block position 0). Logistic regression predictions with a stochastic (blue) and greedy (red) policy are overlaid on probabilities observed for mice (gray). Dark lines show the mean across trials at the same block position and the shading shows the standard error.

The feature weights indicate a propensity of mice to repeat their previous action, as denoted by the positive coefficient on previous choice (hereby denoted by *α*, Fig. 4B).

We tested the fit model on the held-out data to predict the left or right choice of the mice and found that this model, coupled with a stochastic action policy, recapitulated all features of the behavior and achieved comparable log-likelihood estimates on held-out data to those of the nonparametric model (Fig. 4C-F, blue traces; LL = -0.182, Table S3). The “stochastic” policy used here, and in all models below, is a special case where the model selects its port at a rate proportional to the model estimate (see Methods). The stochastic logistic regression captured both the port choice and switching behavior of the mouse as well as possible given the expected confusion matrices (i.e., Δ ≈ 0, Fig. 4C). The model captures both the history dependence of the mouse’s switching behavior, including the apparent stochasticity of conditional switching (Fig. 4D, E). Finally, the model recapitulates the time course over which the block transition perturbs stable port selection and uses increased switch prediction as a mechanism to recover the selection of the high port (Fig. 4F).

These results differ from those of the theoretically motivated model (i.e., HMM) as well as from the same logistic regression model using a deterministic policy (a greedy policy that selects the port with higher log-odds; Fig. 4, red traces). Interestingly, the impact of policy on model performance is most evident when evaluating model fit on switching behavior, with surprisingly subtle effects on the model’s accuracy in predicting left vs. right choice (Fig. 4F). Although the greedy logistic regression captures much of the dynamics of *P*_highchoice_ (Fig. 4F, left), it does so without predicting switching between ports (Fig. 4F, right). These results emphasize the need to explicitly examine switch trials in behavioral modeling.

### Recursive formulation of the reduced logistic regression

Our goal in modeling behavior was to uncover the task features and algorithms that lead to the expressed decision-making strategy. The reduced logistic regression accurately captures the mouse behavior, but requires the weights on each of its features to be learned and the sequence of past choices and rewards to be stored in memory. Furthermore, it requires adapting feature weights when task conditions change such that the animal would essentially need to store multiple look-up tables of feature weights and recall the correct table to perform the task. As such a look-up table-based strategy seems implausible as the foundation of mouse behavior, we inspected the structure of the logistic regression model to determine whether we could achieve similar predictive accuracies with a recursively updated algorithm.

The weights assigned to past choices and rewards were well fit by an exponential curve, with initial magnitude *β* that decays across trials at a rate of *τ* (Fig. 5B). Using this exponential approximation, and approximating the finite sum with an infinite one (since *τ < L*_2_), we can rewrite the log-odds of port selection on the next trial (*ψ*_*t*+1_) as,

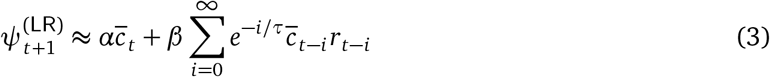

**Figure 5:**
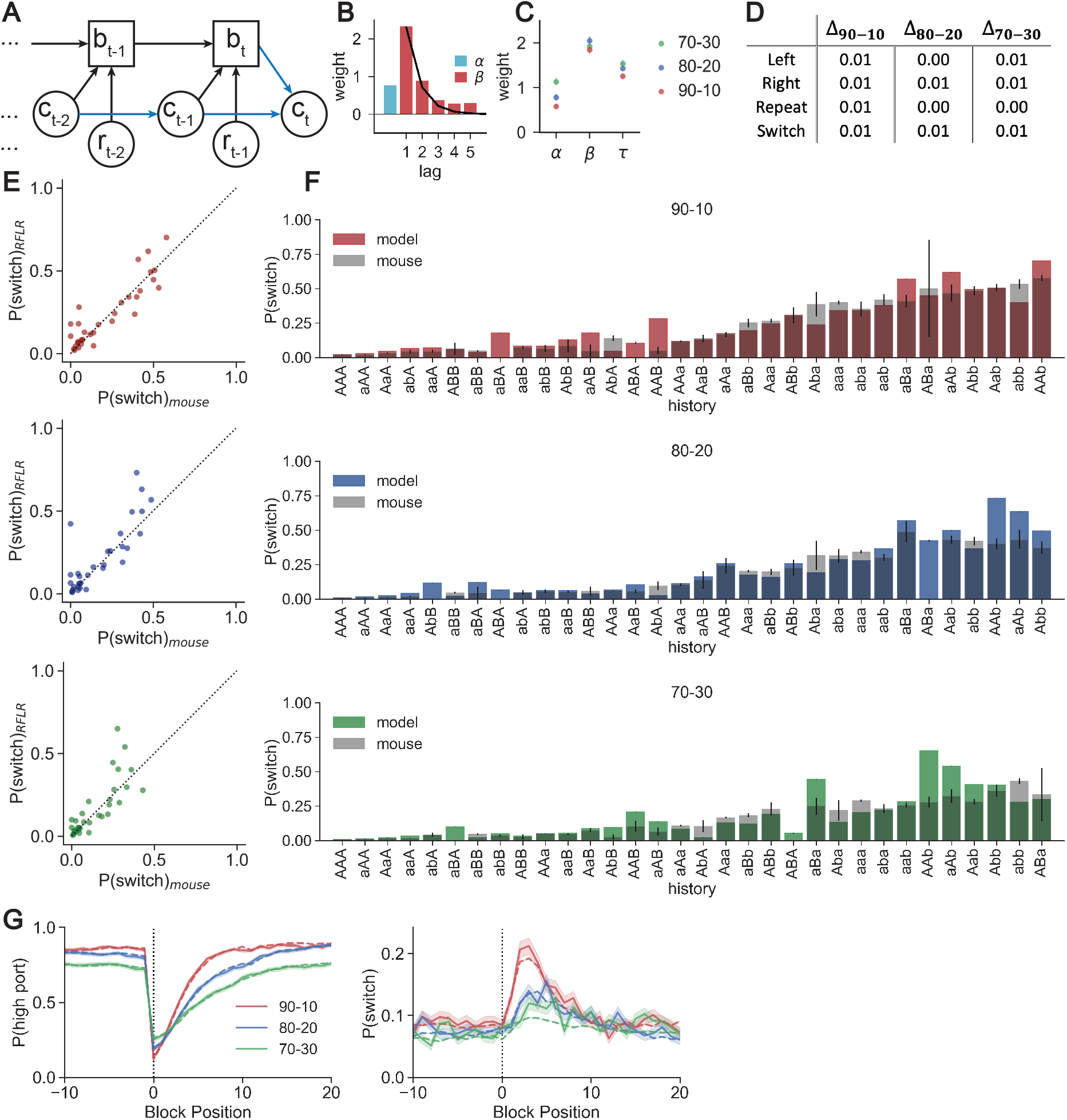
A recursive formulation of the logistic regression recapitulates behavior in multiple reward probability conditions. (A) A recursively formulated logistic regression (RFLR) updates a single state belief (*b*_*t*_) using evidence from recent choice (*c*_*t*− 1_) and reward (*r*_*t*− 1_). The policy (blue) shows an additional contribution on next choice prediction from the previous choice. (B) *β* weights for choice-reward information are described by an exponential function (black curve). (C) Summary of the fit RFLR parameters for data from mice performing in the three sets of reward probability conditions. Each data point shows the mean parameter estimate with error bars indicating the bootstrapped 95% confidence intervals. (D) Δ scores for absolute values of the differences between the RFLR confusion matrices and nonparametric confusion matrix (Fig. 2E) for each action across the three reward probability conditions. (E) Conditional switch probabilities calculated from the RFLR predictions plotted against those of the observed mouse behavior for each set of reward probability conditions (*top*: 90-10 (SSE=0.243), *middle*: 80-20 (SSE=0.417), *bottom*: 70-30 (SSE=0.33)). (F) Conditional switch probabilities predicted by the RFLR (model) across sequences of history length 3 overlaid on those from the mouse data (gray) for the three sets of reward probability conditions. Error bars show binomial standard error for the mouse. (G) *P*_highchoice_ (*left*) and *P*_switch_ (*right*) as a function of trial number surrounding state transition (block position 0) for the three sets of probability conditions. Dashed lines show mean of model predictions, solid lines show mean of true mouse probabilities across trials at the same block position. Shading shows the standard error.

Furthermore, we can compute the infinite sum recursively by observing that,

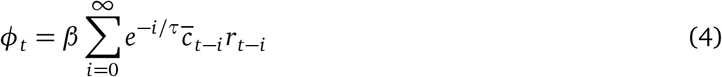

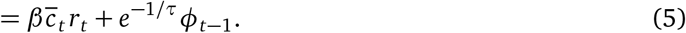

We recognize the resulting form as mathematically analogous to a drift diffusion model [21–23] that decays toward zero with time constant *τ*, but receives additive inputs depending on the most recent choice and whether or not it yielded a reward. The magnitude *β* determines the weight given to incoming evidence. Therefore, our computation of the log-odds can be given as a filtering of choices and rewards biased by *α* toward the most recent choice (Fig. 5A):

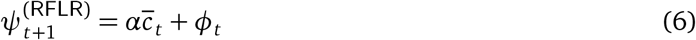

This form of the model offers two advantages over the original logistic regression when considering a potential neural implementation of the algorithm: 1) the exponential representation of choice and reward history captures the behavior using a model with only three parameters (*α, β, τ*), whereas the logistic regression used six, and 2) the recursive definition of this choice-reward representation reduces the memory demands since the model only needs to store the current state estimate (*ϕ*_*t*_), choice (*c*_*t*_), and reward (*r*_*t*_).

We tested this recursively formulated logistic regression (RFLR) on all three sets of reward conditions (i.e., 90-10, 80-20, and 70-30), and found it predicted all features of mouse behavior excellently (Fig. 5D-G, Table S3). Interestingly, the *α* parameter varied the most across reward probability conditions, whereas *β* and *τ* remained relatively constant (Fig. 5C), suggesting that the mechanism by which mice adapt their behavior can be explained by increasing or decreasing their bias toward repeating their previous choice. Notably, *α >* 0 in all contexts, such that there was always some tendency to repeat the previous choice (“stickiness”).

### Relation to reinforcement learning algorithms

Algorithms that use trial-to-trial behavior and outcomes incrementally to build an estimate, rather than requiring recalling the full action history for each new choice, are appealing for decision-making theory. The RFLR resembles another class of such algorithms, namely Q-learning algorithms used in reinforcement learning [19]. Q-learning algorithms use a model-free approach to compute “quality” estimates for each choice available to the agent and recursively update these estimates depending on whether or not a reward is received [19]. Let *Q*_*t*,1_ and *Q*_*t*,0_ denote the quality estimates for the left and right choices, respectively. The recursive updates are,

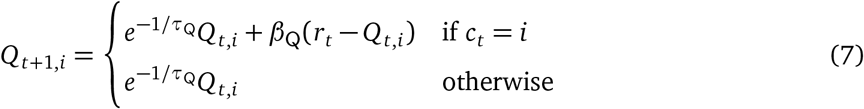

Thus, the quality estimates decay toward zero at a rate determined by the “forgetting” time constant *τ*_Q_, and the chosen port’s quality is updated based on the discrepancy between the received and expected reward and a “learning rate” *β*_Q_. The policy is given by, 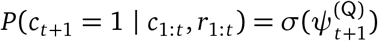, where the log-odds are modeled as

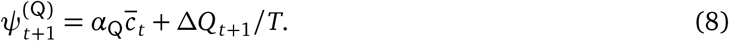

Here, *α*_Q_ is a weight on the previous choice, Δ*Q*_*t*+1_ = *Q*_*t*+1,1_ −*Q*_*t*+1,0_ is the difference in quality estimates, and *T* is a temperature parameter.

Compare these log-odds to those of the RFLR model (eq. 6) and note that Δ*Q*_*t*+1_*/T* is analogous to the recursive quantity *ϕ*_*t*_. It too can be computed recursively:

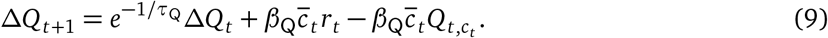

These updates are nearly the same as those for *ϕ*_*t*_ (eq. 5). The only difference is the final term, which depends on the current quality estimate of the chosen port.

In the closely related “forgetting Q-learning” (F-Q) model [24], the final term in eq. 9 disappears. The key difference in the F-Q model is that the updates for the chosen port are replaced with a convex combination of the current estimate and the observed reward, 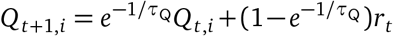, when *c*_*t*_ = *i*. Under this formulation, the dynamics for Δ*Q*_*t*+1_ simplify so that an exact correspondence can be made between the parameters of the RFLR model and those of the F-Q model [30, and see Methods].

As expected, implementing the F-Q model with learning/forgetting, choice-history bias, and temperature values derived from the RFLR model yielded equivalent trial-by-trial choice probabilities and results (LL = -0.182, Table S3). The more flexible Q-learning model did not yield higher performance, and in fact appears to over-fit (LL = -0.185). The result of this analysis provides an algorithmic formulation based on reinforcement learning theory that comprehensively captures mouse choice and switching behavior.

### Returning to the Bayesian agent

The full LR, RFLR, and Q-learning models are similar in both their form and their ability to predict mouse behavior, contrasting the poor performance of the theoretically optimal Bayesian agent. The Bayesian agent uses knowledge of the task structure (i.e. an HMM) to infer the environmental state and guide future action, but it does not capture the tendency of the mice to repeat their last action (i.e., compare graphical representations of the HMM and RFLR in Fig. 3A and 5A).

To gain insight into the differences between these models, we developed a mathematical correspondence for the log-odds computation by the RFLR with that of the Bayesian agent performing inference in an HMM (see Methods). Let 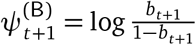 denote the belief state of the Bayesian agent, converted into log-odds. We showed that the recursive belief state calculations for the HMM can be written as,

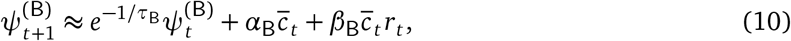

where and (*α*_B_, *β*_B_, *τ*_B_) are determined by the reward and transition probabilities of the HMM (see Methods). These updates are similar to those of the RFLR, allowing us to establish a relationship between the RFLR parameters and the reward and transition probabilities of the HMM.

This analysis revealed two key differences between the RFLR model and the Bayesian agent. First, they differ in how they weigh the preceding choice, 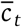. Whereas for all conditions the best-fitting RFLR model yields *α >* 0 (Fig. 5C), the optimal HMM requires *α*_B_ *<* 0 (see derivation in Methods). Intuitively, the RFLR model tends to repeat its previous action, in contrast to the HMM, which makes its selection considering only its posterior belief and independent of any additional choice history bias. Second, the RFLR recursions operate on the weighted sum of past choices and rewards, *ϕ*_*t*_, whereas the HMM recursions operate directly on the log odds, 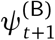. To address this difference, the Bayesian agent needs an additional tendency to repeat its choice immediately after switching ports. We show that the RFLR and the Bayesian agent can be made equivalent by adding a “stickiness” bias,

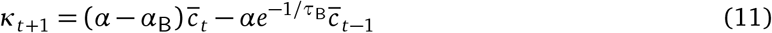

to the Bayesian agent. Then its policy is given by, 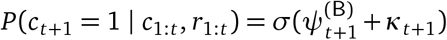 (see Methods). and, by construction, it matches the performance of the descriptive and algorithmic models (Table 3).

### Comparison of behavior of models performing the two-armed bandit task

Analysis of the trial-by-trial log-odds estimates for the RFLR (and accordingly for the sticky HMM, F-Q, and full LR model) reveal asymmetrical use of rewarded vs. unrewarded choice information, whereby rewarded choices provide evidence toward the selected port, but unrewarded choices result in a decay toward *α* (and therefore maintaining a preference for the most recent choice, Fig. 6A). This contrasts the mechanics of the optimal agent, for which unrewarded trials provide evidence toward the alternative port (Fig. 6A). For an optimal agent, an unrewarded choice (or series of unrewarded choices) at the current selection port can flip the sign of belief or log-odds ratio, providing evidence in favor of switching ports (*P*_switch_ *>* 0.5, and even nearing deterministic *P*_switch_), in conflict with the actual mouse behavior.

**Figure 6:**
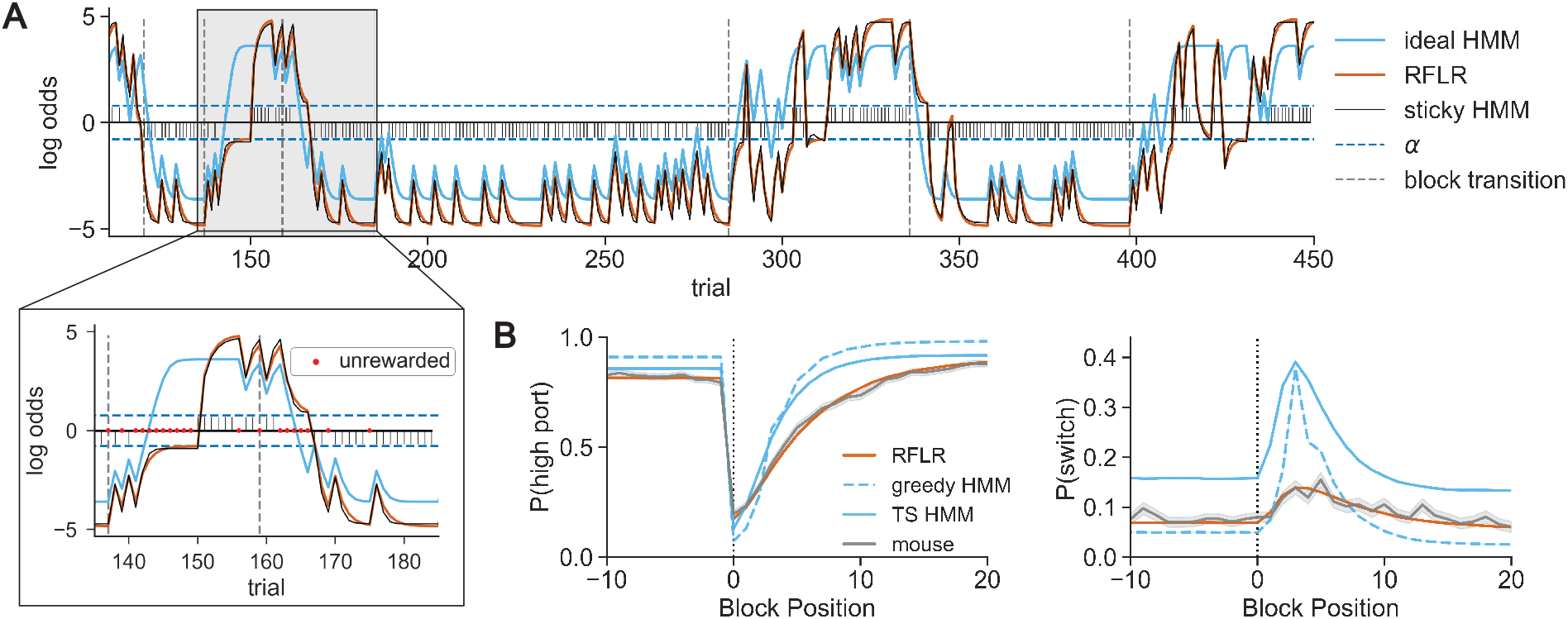
Simulations with a generative recursively formulated logistic regression recapitulate mouse behavior. (A) Representative session depicting equivalent trial-by-trial log-odds computations for the RFLR vs. the sticky HMM (orange vs. black traces). These model estimates contrast the log-odds of the posterior computed by the ideal HMM (light blue), which specifically diverges in prediction updating following unrewarded trials. Stem plot shows the choice*reward interaction that provides action-outcome evidence to the RFLR. Horizontal dashed lines indicate ±*α*, and vertical dashed lines indicate state transitions. Inset: Expanded segment of session with unrewarded trials labeled by red dots. (B) *P*_highchoice_ (*left*) and *P*_switch_ (*right*) as a function of trial number surrounding a state transition (block position 0) in the 80-20 condition for the generative RFLR (orange) and generative ideal HMM (light blue, dashed) and Thompson sampling HMM (TS, solid) overlaid with the observed mouse probabilities (grey). The lines show the means across trials at the same block position and the shadings show the standard errors.

In contrast, for the empirically better-fitting models (LR, RFLR, F-Q, and sticky HMM), the effect of unrewarded trials on the log-odds estimate is to drift towards its choice history bias (i.e. *α*) and, therefore, like the mouse, cause increasingly random port selection. Shifting the port favored by the empirical models requires achieving a reward on the alternative port from the current preference, which causes an update and sign flip in the belief parameter. This suggests that switches under the empirical models rely on the combination of the odds ratio approaching 1 (i.e. log-odds=0) and a stochastic action policy to facilitate random sampling of the low-probability port. It is these stochastic switches — rather than evidence-based switching — that allows the empirical models to update their belief to favor a new action in the future.

The empirical versus ideal models exhibit different bounds on the maximum and minimum trial-to-trial switching probability (see Methods; Fig. 6A). The upper and lower bounds of switching probability in the ideal Bayesian agent are constrained by the odds ratio of the transition probability — even when the model confidently infers the current state of the port reward probabilities, the log-odds are bounded by the probability that the system remains in this state on the next trial. In contrast, the RFLR and sticky HMM reach near-deterministic steady-state behavior (Fig. 6A, Methods). These bounds explain the elevated switch rate produced by Thompson sampling on the HMM belief state, even outside of the block transition (Fig. 6B). Following reward, the belief log-odds of the HMM are further constrained to the product of the odds ratios of the emission probability and transition probability.

### Optimality of behavior

The deviation of the empirical behavior from the theoretically optimal model appears striking when examining trial-by-trial action selection. However, it is unclear that these deviations have a significant cost in terms of the total rewards received. Surprisingly, the expected reward rate of the original Thompson sampling HMM predicting choice from mouse behavior was only marginally better than that actually achieved by the mice (71% vs. 70% trials rewarded in 80-20 sessions, respectively).

To determine whether this was an effect of the suboptimality of the mouse history crippling the HMM performance, we simulated data under the ideal HMM unbounded from mouse history. We initialized an ideal observer model with the true task parameters (*p* = 0.8 and *q* = 0.98) and allowed it to play the game using its own past choices and rewards as history. This model did not perform better, achieving 71% ± 0.4% rewards per session (mean ± s.e.m.). While this version of the HMM uses the optimal inference process for the task, an ideal agent in this task should act greedily on the inferred belief. Indeed, a greedy HMM using the same parameters achieves marginally greater reward rates (Fig. 6B, 74% ± 0.0%), highlighting the significance of the action policy on switching behavior. We compared the performance of these models to simulations run under a generative form of the RFLR using the empirically fit parameters, which achieved 69% ± 0.0% rewards per session (mean ± s.e.m.). Notably, even without the mouse history as input features to guide action selection, the RFLR-generated behavior resembles the characteristic patterns of mouse behavior (Fig. 6B).

We hypothesized that the mice converged to a local maximum or plateau of expected reward within the parameter space in which further optimization of behavior driven by reward rate is challenging. For each of the three reward conditions we held *τ* constant at the corresponding empirically fit value and examined expected reward across the two-dimensional parameter space for varying *α* and *β*. In each, there is a wide plateau over which expected reward stabilizes, and both the *α* and *β* values for the true task parameters under the original HMM and the fit values under the RFLR lie near this plateau (Fig. 7A). For this reason, near-maximal performance can be achieved with a broad range of *α* and *β* values (Fig. 7B).

**Figure 7:**
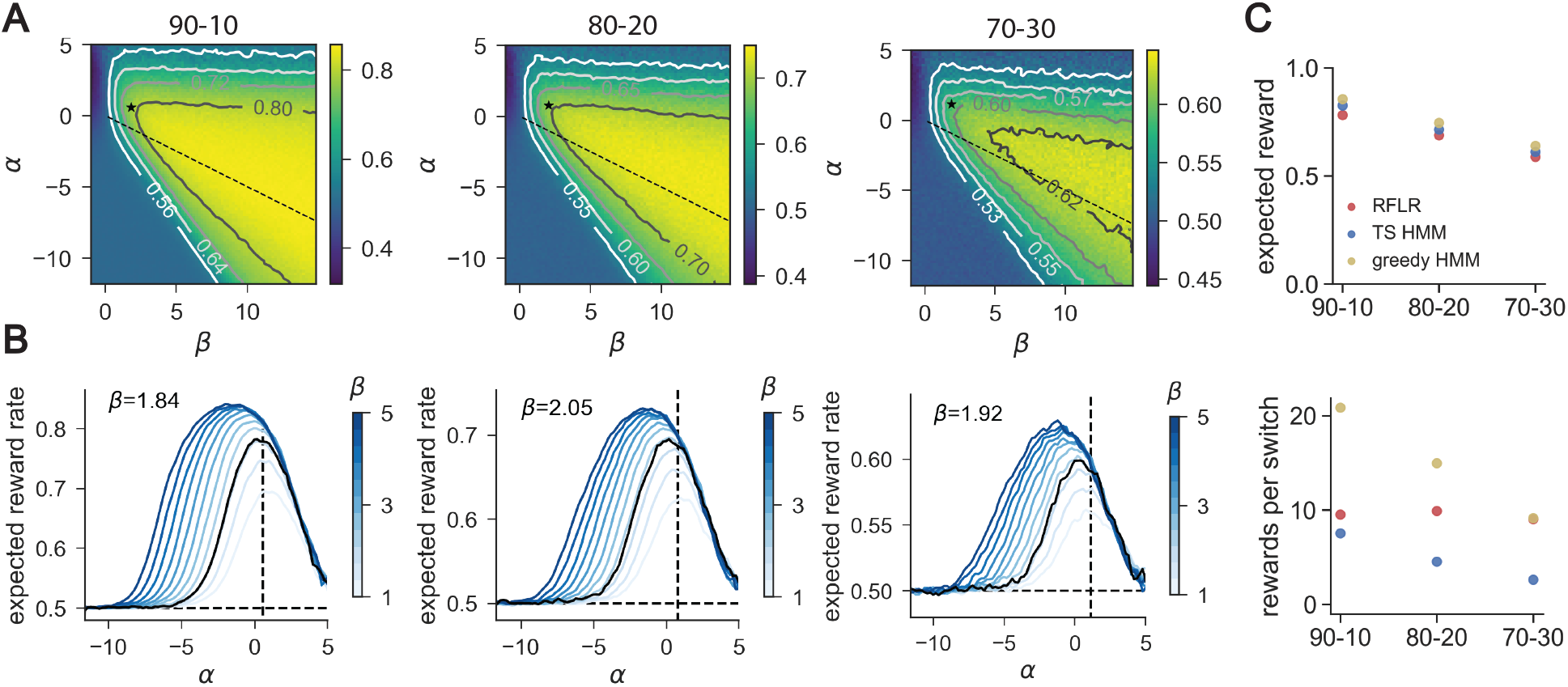
Reward per switch ratios differentiate models and policies that all achieve near-maximal expected reward. (A) Expected reward landscape for the generative RFLR across varying *α* (y-axis) and *β* (x-axis) values with the empirically observed *τ* in each of the three reward conditions (*τ*_90− 10_ = 1.25, *τ*_80− 20_ = 1.43, *τ*_70− 30_ = 1.54). Color bars indicate expected reward rate across simulated trials, and isoclines mark increments above random (0.5). The RFLR-fit *α* and *β* values are depicted at the asterisk (*), and the relative *α* and *β* specified for the HMM lie along the dashed line. (B) Profile of expected reward as a function of *α* for varying values of *β* (color bar, ranging from *β* = 1 to *β* = 5, with fit *β* in black). Expected reward rate at the fit *α* (black vertical dashed line) suggests minimal additional benefit of modulating *β*. (C) *top*: Expected reward in each of the three probability contexts for the generative RFLR using mouse-fit parameters and generative HMM using the true task parameters. HMM performance is shown using either a greedy or stochastic (Thompson sampling, TS) policy. *bottom*: Ratio of rewards to switches for each of the three models across reward probability conditions. Each data point shows the mean across simulated sessions and error bars show standard error but are smaller than the symbol size.

We also considered that the mice may optimize reward relative to a cognitive or physical cost, as opposed to optimizing reward rate at any cost. Specifically, we hypothesized that the stickiness of the empirical models might indicate a preference for the mice not just for reward maximization, but also for efficient collection of reward in terms of behavioral effort, in this case as reflected in the switching rate. Comparing the ratio of rewards to switches, we found that the RFLR achieves twice as many rewards per switch as the Thompson sampling HMM in the 80-20 condition (i.e., an average of 9.95 vs. 4.46 rewards/switch, respectively). Calculating this ratio of rewards per switch for models simulating behavior in each reward context, we find that the RFLR exceeds the original HMM in all three (Fig. 7C). Interestingly, this parallels minimal differences between the models in overall expected reward (Fig. 7C), and so can be attributed to the RFLR’s efficient reduction in switching. However, both of these models are outperformed by the ideal Bayesian agent that uses a greedy policy, indicating that the RFLR’s advantage to maximizing rewards per switch is only valid under the constraint of a stochastic policy. Thus, under the assumption that switching ports bears a cognitive and/or physical cost, and given a tendency for exploration, the objective of the mice may not be exclusively reward maximization, but rather optimizing the tradeoff between reward maximization and cost.

## Discussion

We find that mice performing a two-armed bandit task exhibit switching behavior defined by apparent stochasticity, stickiness, and a representation of action value. These components can be represented in multiple distinct, yet equivalent, models to comprehensively capture both trial-by-trial switching and port choice behaviors. Furthermore, although mouse behavior deviates from that of the ideal Bayesian observer, the expected reward for the empirical models is comparable to that for the ideal agent. Additionally, given a tendency toward exploration, this strategy preserves high reward rates while minimizing trial-by-trial switches via a choice-history bias. Modulating this level of stickiness captures the adaptive response of mice to different reward contexts, offering a parsimonious solution to learning new environmental parameters.

### Switch trials reveal stochasticity in mouse behavior

Many behavioral tasks, including the two-armed bandit, are described as having components of “explore vs. exploit” in which an agent at times exploits existing knowledge and executes an action most likely to lead to reward, whereas at other times it explores the environment by choosing an action with a less certain outcome that reveals information about the environment [9, 12, 14, 31–33].

In such tasks, the trials in which the agent switches actions are the manifestation of exploration and behavioral flexibility (i.e., changes in action due to accumulating information), which are highly informative components of the behavior. Analysis of these trials provides an important insight by revealing the apparently stochastic nature of mouse decision making given recent choice and reward history: although mice enter a regime of nearly deterministic repetition of actions, they do not enter a corresponding regime of deterministic switching (i.e., no accumulation of evidence will consistently push the mouse to switch actions). Thus, even following a series of ‘no reward’ outcomes at a single port, the mouse chooses its next action apparently at random rather than reliably switch selection to the other port.

This understanding propels our selection of a stochastic policy to capture the tendency of the mice to make decisions at a rate proportional to their confidence in those decisions. The policy we fit to behavior balances exploitation of the choice favored by the model estimate and exploration of the alternative choice [9, 16]. The stochasticity we describe is observed under the constraints of our model variables and history length but does not necessarily characterize the decision to switch given an unconstrained model (i.e., given a complete history or access to neural activity). Clearly, at the extreme, the exact sequence of actions and action outcomes expressed by the mouse leading up to a trial late in a session is likely unique (given the exponential growth in sequence possibilities as a function of trial number), and thus it is not possible to determine if the action choice is stochastic given the full history.

### Stochasticity of behavior constrains maximum predictability of behavior by models

There has been a recent push in behavioral studies to account for behavioral events at the resolution of single trials [34]. This is a worthwhile goal, especially in evaluating the predictive performance of behavioral models. However, we found that the stochastic component of behavior given our model assumptions sets bounds on the predictability of different actions. Therefore, we compared the performance of each model against the theoretical probabilities of predicting each action (i.e., expected confusion matrices from the nonparametric model) set by the stochasticity of the mouse behavior on the same type of trial. In the context of exploratory behavior, the method described here or a similar approach to constraining models under the true distribution of the data [33] enables testing of models against realistic boundaries of predictive accuracy.

### Stickiness captures the deviation of mouse behavior from the optimal agent

Interestingly, we find that the model that best recapitulates the mouse behavior, even after the animals have undergone extensive training, does not use the strategy that maximizes reward in this task (the HMM with a greedy policy). Single latent variable HMMs can be implemented in artificial neural networks and therefore, at least in principle, by the brain, so it is unclear why mice do not perform this optimal strategy [35]. An ethological explanation can be proposed from our observation that using the optimal strategy offers only marginal increases in expected reward over the simple recursively formulated logistic regression (or analogous F-Q model; 74% vs. 69% expected reward, respectively). Moreover, given a tendency for exploration or stochasticity, the HMM requires more trials in which the agent switches between ports to achieve equal reward. This hypothesis suggests that constraints imposed by learning the task structure and the asymmetric costs associated with the selection or executions of actions lead the mouse away from the HMM implementation. Additionally, it brings up an interesting question as to whether mice have an innate tendency for exploration in environments with uncertainty [36–39].

We found that a descriptive model, the logistic regression, offered a better fit to mouse behavior than the ideal observer model. This finding is consistent with recent data-driven modeling of rodents behaving in a similar two-armed bandit task in which the reward contingencies drifted continuously from one trial to the next [13]. In that case, the logistic regression coefficients exhibited a similar exponential decay, placing the greatest weight on the most recent actions and outcomes. Given that both tasks involve reward contingencies with Markovian dynamics, we suspect that a similar connection could be made to a theoretically motivated ideal observer model performing inference in an HMM and minimizing switching.

In our analysis, the differences between the ideal observer and the data-driven models were explained by an additional influence of past choice on future choice. We accounted for this by building a sticky HMM, which, by construction, produced equivalent trial-by-trial log-odds predictions as the RFLR. Stickiness has been reported in analyses of behavior across tasks and species, and is also called perseveration, choice history bias, and the law of exercise [13, 23, 24, 39–43]. This bias to repeat previous actions offers a parsimonious mechanism for adapting an existing action policy to novel environmental conditions: we found that, in the face of changing reward probability conditions, mice minimally updated the weight assigned to incoming evidence and the time constant of memory decay (*β* and *τ*, respectively), but instead modulated their behavior by increasing or decreasing their level of perseveration. In the mathematically equivalent F-Q formulation of reinforcement learning, this change is conserved as the *α* parameter is directly derived from the RFLR. This behavioral adaptation, represented largely by a single parameter, comes at low cost to the animal in terms of expected reward, and therefore may be an efficient strategy for minimizing effort necessary to learn new behavioral strategies [43–45].

### Implications for the neural mechanism

One of the goals of this study was to increase our understanding of decision making in order to guide future interrogation of circuit function and the neural underpinnings of behavior. However, the specific algorithms that we found best fit the mouse behavior may or may not be directly implemented in the brain. The demonstration that multiple distinct algorithms can similarly model behavior underscores this point and draws our focus to the features that are shared by the models. We hypothesize that whatever algorithm the brain relies on for this task, if it is deterministic then it is combined with a stochastic action policy to produce the behavior we observe. (Of course, a policy that appears stochastic behaviorally can have deterministic neural origins.) Recently developed statistical methods offer new means of determining if and how the features of these behavioral models are encoded in neural activity [46].

Each of the models that successfully recapitulates mouse behavior relies on an interaction between choice and reward, consistent with previous accounts of action value encoding in brain regions such as striatum and medial prefrontal cortex [5, 6, 10, 11, 47]. The action value representation in our empirical models notably treats evidence from rewarded and unrewarded trials asymmetrically. This asymmetry has been previously reported in analysis of mouse evidence accumulation [16], and it contrasts the behavior of the ideal agent that uses unrewarded trials as evidence in favor of the alternative option. Investigating whether, and at what level of processing, a corresponding asymmetry exists in the neural representation of reward will be important for understanding the nature of reinforcement in learning.

Furthermore, past work has hypothesized that recursive algorithms that compress information over a sequence of trials to a small number of variables are more neurally plausible [31]. Here, we show that some recursive algorithms (i.e., original HMM) struggle to explain switching behavior, whereas non-recursive models (i.e., logistic regression) perform well. This poses a potential challenge to this hypothesis. However, we were able to derive alternative recursive algorithms (i.e., RFLR, F-Q model, and sticky HMM) that do accurately explain behavior.

Is there a way to disambiguate these models if they all produce the same behavior? Previous work has shown that model-based and model-free methods may exist in parallel, but can be distinguished through measurements of neural activity [48]. Behaviorally, model-based representations may offer an advantage for overall performance accuracy, but as has been shown here and previously, the magnitude of this difference in accuracy is task-dependent [49]. Importantly, model-based implementations come at a cost for cognitive demands [49], so this tradeoff between demand and reward may favor model-based methods in some contexts but not others. Additional contextual features may make the implementation of particular models favorable, such as environments where action outcomes are not symmetric or inter-dependent. In this case, the ability to separately approximate the value of each action, as in Q-learning models, could be beneficial. Furthermore, in a lateralized task, this could allow for lateralized representations across hemispheres.

More complex behavioral models might predict mouse behavior more accurately than those discussed here. For example, recent work has shown that mouse behavior in similar tasks is well-described by drifting or discretely switching policies, suggesting that animal behavior is guided by time-varying internal states [50, 51]. This raises an interesting and perhaps confusing point: as we seek to understand animal behavior, we must simultaneously infer an animal’s internal state as well as that animal’s inferences about the external state of the world. Probabilistic models that bridge descriptive, algorithmic, and theoretically-guided characterizations offer a route to resolving these complexities of animal behavior.

## Materials and Methods

### Behavior apparatus

The arena for the two-armed bandit task was inspired by previous work [6]. Behavior experiments were conducted in 4.9” x 6” custom acrylic chambers. Each chamber contained three nose ports with an infrared-beam sensor (Digi-Key, 365-1769-ND) to detect entry of the snout into the port. A colored LED was positioned above each port. For the two side ports, water was delivered in 2.5 µL increments via stainless steel tubes controlled by solenoids (The Lee Co, LHQA0531220H). The timing of task events was controlled by a microcontroller (Arduino) and custom software (MATLAB). Plans for an updated version of the behavioral system, including the most recent hardware and software, are available online: https://edspace.american.edu/openbehavior/project/2abt/ and https://github.com/bernardosabatinilab/two-armed-bandit-task

### Behavior task

Wild-type mice (C56BL/6N from Charles River and bred in house) aged 6-10 weeks were water restricted to 1-2 mL per day prior to training and maintained at >80% of full body weight. While performing the task, mice moved freely in the chamber. Activation of an LED above the center port indicated that the mouse could initiate a trial by nose poking into the center port. Doing so activated LEDs above the two side ports, prompting the mouse to choose to nose poke to the right or left. The mouse had 2 s to make its selection. Following side port entry, the computer determined whether or not to deliver a water reward according to the corresponding port reward probability and the result of pseudo-random number generation. Withdrawal from the side port ended the trial and started an inter-trial interval (ITI). The 1 s ITI followed selection, during which time the system assigned the reward probabilities for the next trial according to a Markov decision process (0.98 probability that high and low port assignments remained the same, 0.02 probability the assignments reversed). After the 1 s minimum ITI, the center port LED turned on and the mouse was permitted to initiate the next trial (with no upper limit to trial initiation time). The duration of each behavior session was 40 minutes, over which the mouse typically earned >350 rewards. All training sessions were conducted in the dark or under red-light conditions.

### Conditional switch probabilities

To concisely represent the history preceding each trial, we defined a code that captures both action (relative choice direction) and the outcome of that action (reward or no reward): the letter (a/A vs b/B) denotes the action and the case (lower vs. upper) denotes the outcome with upper case indicating a rewarded trial. We define the first choice direction of the sequence as “A”, so that, depending on reward outcome, choices in this direction are also labeled “A/a” whereas those in the other direction are labeled “B/b.” This code was used to build “words” (e.g. Aab) that fully specify the action and outcome histories for a given length on which switch probabilities were conditioned.

### Models of mouse behavior

All behavior models were trained on 70% of sessions and tested on the remaining held-out data. For models predicting mouse choice on previous mouse behavior (Fig. 3-5), model predictions were taken as the mean across 1000 repetitions on bootstrapped test data to acquire representative estimates of choice and switch probabilities.

We simulated mouse behavior by implementing the Bayesian agent and RFLR as generative models. We simulated the task with the location of the high rewarding port (*p* = 0.8) determined by a Markovian process with a transition probability of 0.02, and preserved the session structure that the mice experienced, such that the number of trials in each session was drawn from a distribution based on the mouse behavior. Each model was given the same set of sessions, and played until a simulated dataset the same size as the mouse dataset was generated. We ran this simulation for 1000 repetitions to create the averaged performance presented in Figure 6.

For the Bayesian agent, we used the ideal observer given the true task parameters, and after random initialization for the first choice allowed the model to recursively update its belief given its own actions and associated outcomes to guide future choices. We generated behavior from an HMM Thompson sampling on its belief to correspond with the stochastic policy of the RFLR, as well as acting greedily on its belief to represent the ideal observer (Fig. 6B, 7C). For the RFLR, the model played using the fit parameters of the mouse. The expected reward landscape was calculated by performing a parameter grid search with this simulation.

Complete mathematical details are given in the supplementary information text.

## Acknowledgments

We thank Linda Wilbrecht and members of her lab for advice in how to implement the two-armed bandit task. We thank the Harvard Medical School Research Instrumentation Core for help in designing and implementing the necessary hardware and software. We thank members of the Sabatini and Linderman labs for helpful advice and comments on the study and manuscript. We thank Julie Locantore for assistance in training mice. This work was funded by grants to BS and SL from the Brain Initiative (NINDS, U19NS113201, a.k.a Team Dope) and the Simons Collaboration on the Global Brain. Predoctoral fellowships from the NSF graduate research fellowship program supported CB and from the Canadian Institutes of Health Research supported SN. The freely-moving 2ABT behavior was developed under NS046579-13.

