## Supplemental Information for "Mice exhibit stochastic and efficient action switching during probabilistic decision making"

### Mathematical details

This section describes the mathematical details of the Bayesian agents, the logistic regression model and its recursive formulation, the Q-learning algorithm and the special case of the F-Q algorithm, and the derivation of the sticky Bayesian model that matches the empirical mouse behavior.

#### Bayesian models of mouse port choice behavior

Bayesian agents compute a posterior belief over the environmental state given past choices and rewards. For this task, computing the posterior requires inference in an HMM to obtain the belief state, $b_{t+1} = P(z_{t+1} = 1 \mid c_{1:t}, r_{1:t})$, that on the next trial $t$ the environment is in the "left" state ($z_{t+1} = 1$). This belief state can be updated recursively. To compute the next belief state $b_{t+1}$ given the current belief state, we first condition on the current choice and reward and then normalize to obtain,

$$b_t^{(c)} = \frac{P(r_t \mid z_t = 1, c_t)\, b_t}{P(r_t \mid z_t = 0, c_t)\,(1 - b_t) + P(r_t \mid z_t = 1, c_t)\, b_t}. \tag{1}$$

Then we evaluate the probability of the next environmental state given the conditioned beliefs,

$$b_{t+1} = P(z_{t+1} = 1 \mid z_t = 0)\,(1 - b_t^{(c)}) + P(z_{t+1} = 1 \mid z_t = 1)\, b_t^{(c)}. \tag{2}$$

These simple belief state recursions are possible because of the Markovian nature of the model.

We implemented the HMM inference using Python code available on GitHub (`https://github.com/lindermanlab/ssm`). For the Bayesian agents, we built a model with two discrete latent states and access to the true transition and emission matrices. For example, for the version of the task where P(reward | high-choice) = 0.8, the emission probabilities $P(r_t \mid z_t, c_t) \in \{0.8, 0.2\}$ and transition probabilities $P(z_{t+1} \mid z_t) \in \{0.98, 0.02\}$. To test the HMM's performance at predicting mouse choice, we fed the model action and outcome data from sequences of trials. For each session, the model was initialized with equal priors for the right and left state, after which the model iteratively updated its belief by advancing through the trial sequence. $b_{t+1}$ has upper and lower bounds constrained by the nonstationary dynamics of the latent state, captured by the predictive probability of $P(z_{t+1} \mid z_t)$.

[1] Howard Hughes Medical Institute, Department of Neurobiology, Harvard Medical School, Boston, MA
[2] Department of Statistics and Wu Tsai Neurosciences Institute, Stanford University, Stanford, CA
* ``
† ``

The posterior estimate for each trial was passed through a softmax action policy to make a prediction of mouse behavior,

$$P(c_{t+1} = 1 \mid c_{1:t}, r_{1:t}) = \pi(b_{t+1}) = \sigma\big(\sigma^{-1}(b_{t+1})/T\big), \tag{3}$$

where $\sigma(x) = \frac{1}{1+e^{-x}}$ is the logistic function, $\sigma^{-1}(x) = \log\frac{x}{1-x}$ is its inverse, the logit function, and $T \geq 0$ is a temperature parameter. As $T$ goes to zero, this reduces to the greedy policy in which $\pi(b_{t+1}) = 1$ if $b_{t+1} \geq 0.5$ and $\pi(b_{t+1}) = 0$ otherwise. When $T = 1$ the softmax policy is equivalent to Thompson sampling, where $\pi(b_{t+1}) = b_{t+1}$. For intermediate temperatures, this policy can interpolate between the two.

### Bayesian agent fit to mouse behavior

We considered that the true transition and emission probabilities are not known to the mouse. To optimize the fit of our Bayesian agent to the mouse's behavior, we ran a grid search over transition probabilities, emission probabilities, and softmax policy temperatures on bootstrapped training data and calculated the log likelihood of the data given the model using each parameter set. We maximized this function to select the parameters used in our "fit HMM" and evaluated on bootstrapped test dataset (30% of data) as presented.

### Logistic regression

We compute the conditional probability of choice given data from previous choices and rewards using a logistic regression to compute the log-odds, $\psi$:

$$P(c_{t+1} = 1 \mid c_{1:t}, r_{1:t}) = \sigma(\psi^{(\mathrm{LR})}_{t+1}) \tag{4}$$

The full logistic regression model uses the weighted linear combination of the action (choice), reward outcome, and action-outcome interaction history from previous trials to calculate the log-odds of mouse choice for the next trial:

$$\psi^{(\mathrm{LR})}_{t+1} = \sum_{i=0}^{L_1} \alpha_i \bar{c}_{t-i} + \sum_{i=0}^{L_2} \beta_i \bar{c}_{t-i} r_{t-i} \sum_{i=0}^{L_3} \gamma_i r_{t-i} + \delta \tag{5}$$

where $\alpha$, $\beta$, and $\gamma$ represent the weights on input features for choice, encoding of choice-reward interaction, and reward across trials back to $L_1$, $L_2$, and $L_3$, respectively. For the features, $\bar{c}_{t-i}$ represents whether the mouse made a choice to the left or right (1 or -1 respectively), $r_{t-i}$ represents whether or not the mouse received a reward (1 or 0, respectively), $\bar{c}_{t-i} r_{t-i}$ the interaction between choice and reward (1 when rewarded left, -1 when rewarded right, 0 otherwise) on the i-th trial back, and $\delta$ represents the overall port bias. To fit the model, we split our data into training and testing datasets. We used cross validation to fit the hyperparameters $L_1$, $L_2$, and $L_3$ as 1, 5, and 0 respectively, to arrive at the reduced form of the model presented in the text and Figure 4:

$$\psi^{(\mathrm{LR})}_{t+1} = \alpha \bar{c}_t + \sum_{i=0}^{5} \beta_i \bar{c}_{t-i} r_{t-i} \tag{6}$$

### Recursive formulation of the empirical logistic regression

We use the exponential function, $\beta_i = \beta e^{-i/\tau}$ to approximate the weights on the encoded choice-reward history. Substituting this approximation in the reduced logistic regression, we compute the log-odds as:

$$\psi^{(\mathrm{LR})}_{t+1} \approx \alpha \bar{c}_t + \beta \sum_{i=0}^{\infty} e^{-i/\tau} \bar{c}_{t-i} r_{t-i} \tag{7}$$

from which we define the recursive quantity, $\phi_t$,

$$\phi_t = \beta \sum_{i=0}^{\infty} e^{-i/\tau} \bar{c}_{t-i} r_{t-i} \tag{8}$$

$$= \beta \bar{c}_t r_t + e^{-1/\tau} \beta \sum_{i=0}^{\infty} e^{-i/\tau} \bar{c}_{t-1-i} r_{t-1-i} \tag{9}$$

$$= \beta \bar{c}_t r_t + e^{-1/\tau} \phi_{t-1}. \tag{10}$$

Thus, we define the log-odds for the RFLR model as:

$$\psi_{t+1}^{(\mathsf{RFLR})} = \alpha \bar{c}_t + \phi_t \tag{11}$$

Finally, note that we can obtain a recursive formula for the log-odds directly by substituting in the definition of $\phi_t$:

$$\psi_{t+1}^{(\mathsf{RFLR})} = \alpha \bar{c}_t + \beta \bar{c}_t r_t + e^{-1/\tau} \phi_{t-1} \tag{12}$$

$$= e^{-1/\tau} \psi_t^{(\mathsf{RFLR})} + \alpha \bar{c}_t - e^{-1/\tau} \alpha \bar{c}_{t-1} + \beta \bar{c}_t r_t. \tag{13}$$

This formulation will be useful when comparing to the HMM.

We fit the free parameters $\alpha$, $\beta$, and $\tau$ using stochastic gradient descent (reparameterizing $\tau \to \log \tau$ to be unconstrained) on the training set and estimated parameter error with bootstrapped confidence intervals.

### Equivalence of the RFLR and F-Q-learning models

As described previously [30], the "forgetting Q learning" (F-Q) model assumes slightly different updates for the quality estimates,

$$Q_{t+1,i} = \begin{cases} e^{-1/\tau_{\mathsf{Q}}} Q_{t,i} + (1 - e^{-1/\tau_{\mathsf{Q}}}) r_t & \text{if } c_t = i \\ e^{-1/\tau_{\mathsf{Q}}} Q_{t,i} & \text{otherwise} \end{cases} \tag{14}$$

and the policy is determined by,

$$P(c_{t+1} = 1 \mid c_{1:t}, r_{1:t}) = \sigma(\psi_{t+1}^{(\mathsf{Q})}) \tag{15}$$

$$\psi_{t+}^{(\mathsf{Q})} = \alpha_{\mathsf{Q}} \bar{c}_t + \Delta Q_{t+1}/T, \tag{16}$$

where $\Delta Q_{t+1} = Q_{t+1,1} - Q_{t+1,0}$, $T \geq 0$ is a temperature parameter, and $\tau_{\mathsf{Q}} \geq 0$ is a time constant that determines both the "learning rate" and the "forgetting rate". In the F-Q model, the difference in quality estimates evolves as,

$$\Delta Q_{t+1} = e^{-1/\tau_{\mathsf{Q}}} \Delta Q_t + (1 - e^{-1/\tau_{\mathsf{Q}}}) \bar{c}_t r_t \tag{17}$$

The temperature-rescaled difference in quality estimates, $\Delta Q_t / T$, is analogous to the recursive quantity $\phi_t$ in the RFLR model. To equate these two models, we set the RFLR parameters to $\alpha = \alpha_{\mathsf{Q}}$, $\beta = (1 - e^{-1/\tau_{\mathsf{Q}}})/T$, and $\tau = \tau_{\mathsf{Q}}$.

### Mathematical correspondence between HMM and RFLR

The HMM is characterized by the belief state $b_{t+1} = P(z_{t+1} = 1 \mid c_{1:t}, r_{1:t})$ representing the distribution over the latent state given preceding choices and rewards. Let

$$\psi_{t+1}^{(\mathsf{B})} = \sigma^{-1}(b_{t+1}) = \log \frac{b_{t+1}}{1 - b_{t+1}} \tag{18}$$

denote the log-odds ratio of the belief state at trial $t+1$. Under a Thompson sampling policy, we have $P(c_{t+1} = 1 \mid c_{1:t}, r_{1:t}) = \sigma(\psi_{t+1}^{(\mathrm{B})})$.

Eq. 2 gives a recursive update for the belief given new information obtained on each trial. We can rewrite this recursion in terms of the log-odds instead. We find that,

$$\psi_{t+1}^{(\mathrm{B})} = f(\psi_t^{(\mathrm{B,c})}) \tag{19}$$

where

$$\psi_t^{(\mathrm{B,c})} = 2\sigma^{-1}(p)\, r_t \bar{c}_t - \sigma^{-1}(p)\, \bar{c}_t + \psi_{t-1}^{(\mathrm{B})} \tag{20}$$

and

$$f(\psi_t^{(\mathrm{B,c})}) = -\sigma^{-1}(q) + \log \frac{1 + e^{\psi_t^{(\mathrm{B,c})} + \sigma^{-1}(q)}}{1 + e^{\psi_t^{(\mathrm{B,c})} - \sigma^{-1}(q)}} \tag{21}$$

Here, $q \in [\frac{1}{2}, 1)$ is the probability that the system state remains the same and $p \in [\frac{1}{2}, 1)$ is the probability of receiving a reward upon choosing the correct port. (Due to the symmetric design of the experiment, $p$ is also the probability of not receiving a reward upon choosing the incorrect port.)

Though these equations may look rather complicated, they have many intuitive properties. First, the log-odds recursions split into a "conditioning" step in which current log-odds $\psi_t^{(\mathrm{B})}$ are updated with new information from the current choice $\bar{c}_t$ and reward $r_t$ to obtain $\psi_t^{(\mathrm{B,c})}$. This step depends on the same features as the logistic regression model presented above, and the coefficients are functions of the reward probability $p$. Second, the "prediction" step passes the conditioned log-odds through a nonlinear transformation $f$ to obtain log-odds for the next trial. This nonlinear function saturates at $\pm\sigma^{-1}(q)$, implying that the log-odds cannot exceed the log-odds of the transition probability. This makes sense since, even if the mouse knew the state at trial $t$, there is always probability $1-q$ that it will change on the next trial. Moreover, the nonlinearity is steepest when there is substantial uncertainty ($\psi_t^{(\mathrm{B,c})} \approx 0$). In that regime, a rewarded choice has a large influence, whereas when the mouse is already quite certain, one more rewarded choice won't change the log-odds by much.

To make the correspondence between this nonlinear function and the linear updates of the RFLR, we make a first-order Taylor approximation to the HMM prediction function around zero,

$$f(\psi_t^{(\mathrm{B,c})}) \approx f(0) + f'(0)\psi_t^{(\mathrm{B,c})} = (2q-1)\psi_t^{(\mathrm{B,c})}. \tag{22}$$

Substituting the definition of $\psi_t^{(\mathrm{B,c})}$ froim eq. 20 yields a form of the log-odds update that resembles the RFLR update,

$$\psi_{t+1}^{(\mathrm{B})} \approx (2q-1)\big[2\sigma^{-1}(p)\,\bar{c}_t r_t - \sigma^{-1}(p)\,\bar{c}_t + \psi_t^{(\mathrm{B})}\big] \tag{23}$$

$$= e^{-1/\tau_{\mathrm{B}}}\psi_t^{(\mathrm{B})} + \alpha_{\mathrm{B}}\,\bar{c}_t + \beta_{\mathrm{B}}\bar{c}_t r_t \tag{24}$$

where

$$e^{-1/\tau_{\mathrm{B}}} = 2q-1 \tag{25}$$

$$\alpha_{\mathrm{B}} = -(2q-1)\sigma^{-1}(p) \tag{26}$$

$$\beta_{\mathrm{B}} = 2(2q-1)\sigma^{-1}(p) \tag{27}$$

This form highlights the connection to the RFLR model. As the HMM has only two free parameters, $p$ and $q$, the weights are constrained so that $\alpha_{\mathrm{B}} = -\beta_{\mathrm{B}}/2$. Moreover, since $p, q \in [0.5, 1]$, we can see that $\beta_{\mathrm{B}} \geq 0$ and $\alpha_{\mathrm{B}} \leq 0$. This is a key difference from the RFLR model, where $\alpha$ was positive. Likewise, comparing eq. 13 to eq. 23, the other difference is that the HMM updates do not depend on $\bar{c}_{t-1}$.

### Reconciling the HMM and RFLR by adding "stickiness"

The discrepancy between the HMM and the RFLR can be fixed via a simple, "sticky" modification to the HMM to favor previous choices. Suppose we have a RFLR model with parameters $\alpha$, $\beta$, and $\tau$. We map that to a sticky HMM by setting $p$ and $q$ such that $\tau_{\mathsf{B}} = \tau$ and $\beta_{\mathsf{B}} = \beta$. Once these are set, $\alpha_{\mathsf{B}}$ is determined by eq. 26. Then, we add an additional term, $\kappa_{t+1}$, to the HMM log-odds recursions to account for the difference between eq. 13 and eq. 23:

$$\kappa_{t+1} = \psi_{t+1}^{(\mathsf{RFLR})} - \psi_{t+1}^{(\mathsf{B})} = (\alpha - \alpha_{\mathsf{B}})\bar{c}_t - e^{-1/\tau_{\mathsf{B}}}\alpha\bar{c}_{t-1} \tag{28}$$

Alternatively, we can write this expression as,

$$\kappa_{t+1} = [(1 - e^{-1/\tau_{\mathsf{B}}})\alpha - \alpha_{\mathsf{B}}]\bar{c}_t + 2e^{-1/\tau_{\mathsf{B}}}\alpha\bar{c}_t s_t \tag{29}$$

where we have introduced the binary variable $s_t$ to indicate whether or not the mouse switched ports on the previous trial. (That is, $s_t = 1$ if $c_t \neq c_{t-1}$ and $s_t = 0$ if $c_t = c_{t-1}$.) The positive weight on $\bar{c}_t s_t$ is additional reinforcement for exploring a new port, and it indicates that mice are more likely to return to a port immediately after switching to it.

Under the sticky HMM, the policy is,

$$P(c_{t+1} = 1 \mid c_{1:t}, r_{1:t}) = \sigma(\psi_{t+1}^{(\mathsf{B})} + \kappa_{t+1}), \tag{30}$$

which matches the policy of the RFLR model by construction.

### Supplementary Figures

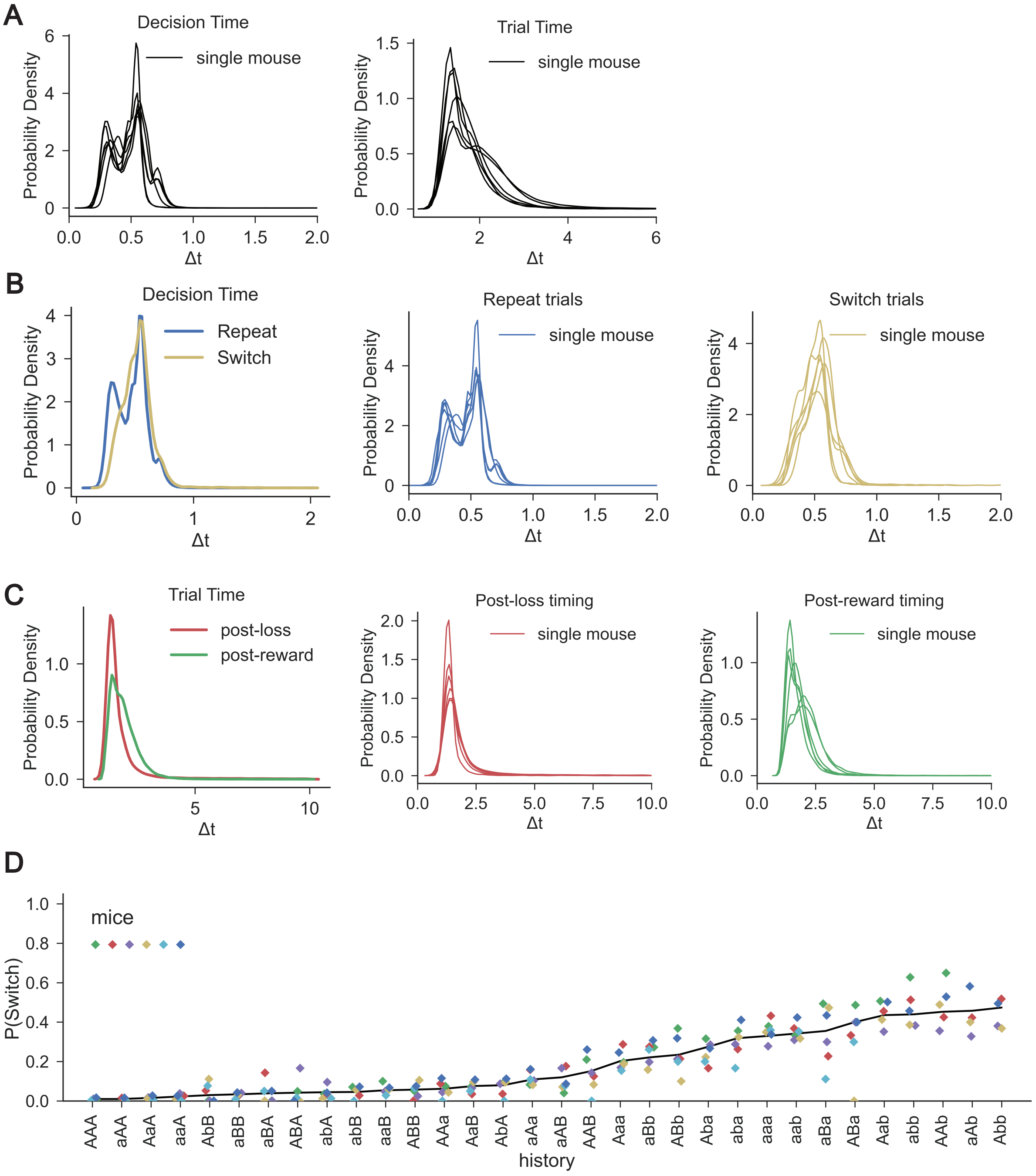


**Figure 1:** Individual mouse behavior and decision times. (A) Probability density distributions of trial times (in seconds) for each mouse (thin line). *left*: Distribution of time from center port to side port (decision time), *right*: distribution of time from center port to center port (trial initiation to next trial initiation). The most extreme trials ($\Delta t > 10s$, comprising <1% trials) were excluded. Probability densities integrate to 1 across the continuous distribution of durations. (B) *left*: Probability density distributions of decision times (center port to side port) for trials in which the mouse switched ports vs. those in which the mouse repeated its decision at the same port. *middle*: Distributions for individual mice on "repeat" trials, *right*: distributions for individual mice on "switch" trials. (C) *left*: Probability density distributions of trial duration (center port to center port, including intertrial interval) following reward vs. following no reward. middle: Distributions for individual mice following no reward, *right*: distributions for individual mice following reward. (D) Conditional switch probabilities by mouse for each action-outcome trial sequence of history length 3. Each symbol indicates the mean switch probability following the corresponding action-outcome history across all trials for each mouse. History sequences are sorted by the aggregate conditional switch probabilities of all mice (black line). Data is from the 80-20 condition.

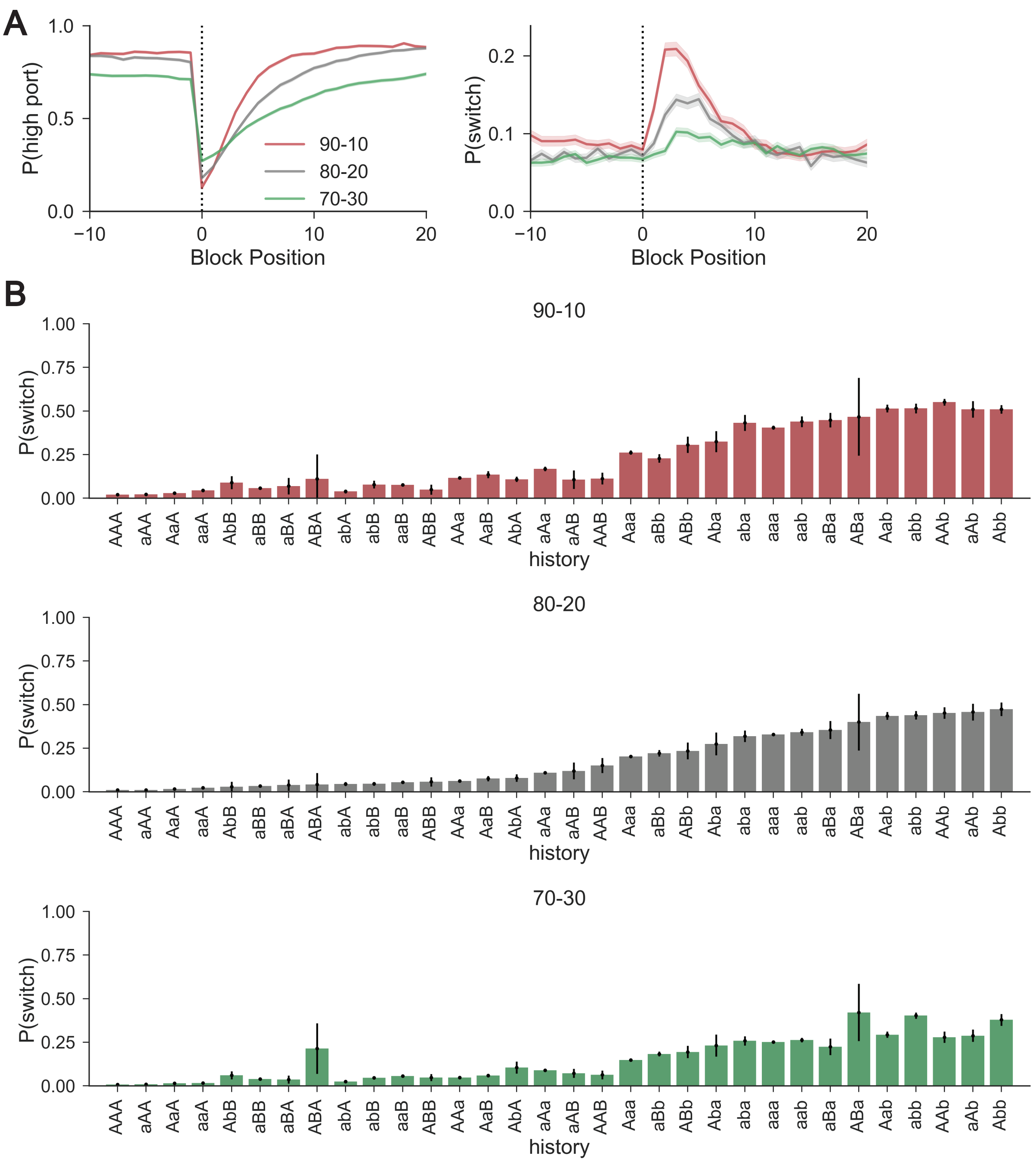


**Figure 2:** Mouse behavior across different reward contexts. (A) $P_{high-choice}$ (*left*) and $P_{switch}$ (*right*) as a function of trial number surrounding the state transition (block position 0) for each reward probability set. Dark lines show the mean across trials at the same block position and the shading shows the standard error. (B) Conditional switch probabilities for mice performing the task with $p = 0.9$ (*top*), $p = 0.8$ (*middle*, as in Figure 2D), and $p = 0.7$ (*bottom*). For each reward probability set, $P(reward|low-choice) = 1 - p$.

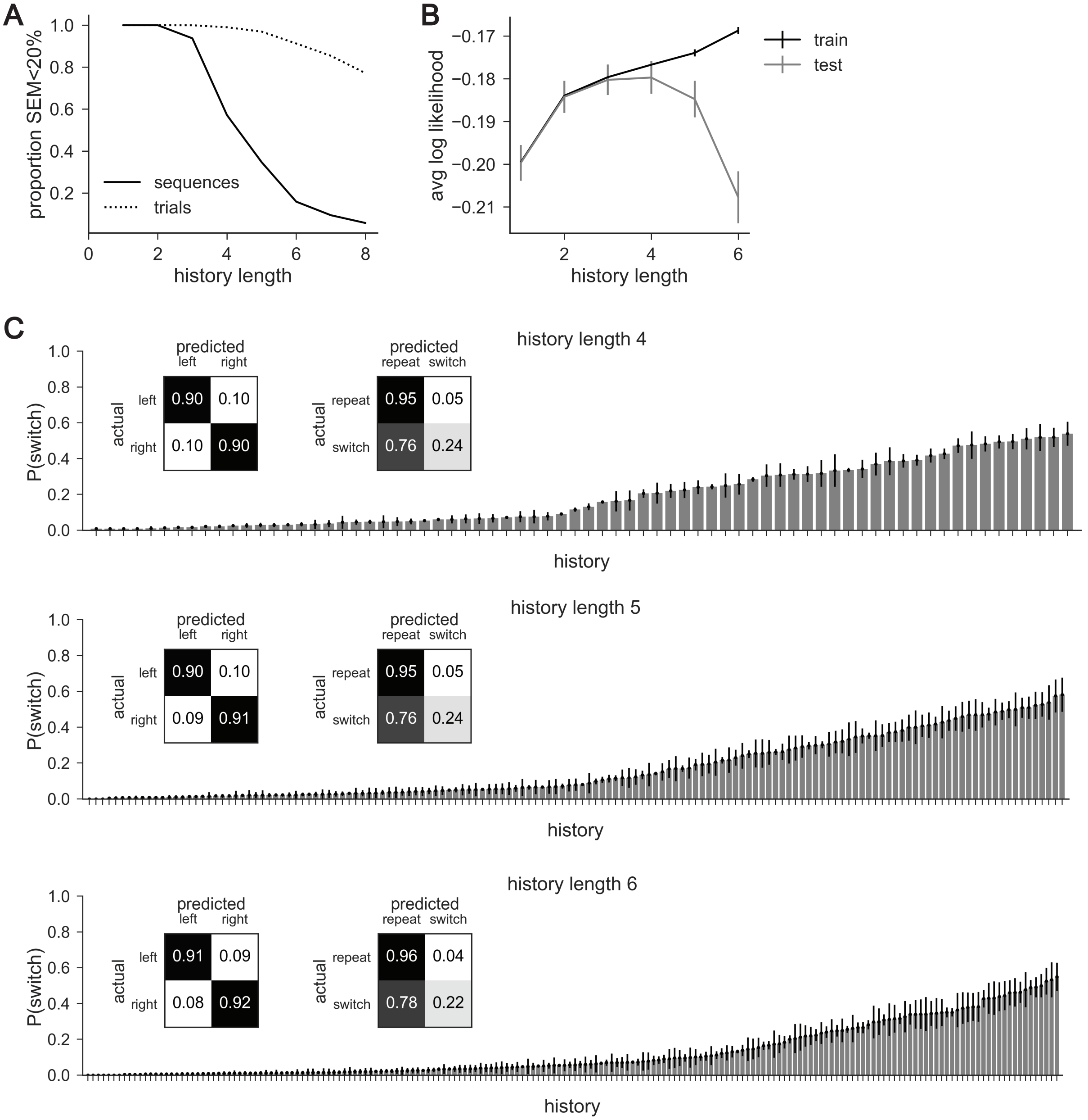


**Figure 3:** Contribution of longer trial histories to conditional switch probabilities. (A) Proportion of expressed action and outcome history sequences with standard error (SEM) less than 20% (solid line), and the corresponding proportion of trials (dashed line) as a function of increasing history length. (B) Average log-likelihood estimates for the empirical nonparametric policies on training and testing data, using 5-fold cross validation. Error bars show standard error of the estimates. (C) Conditional switch probabilities given action-outcome trial sequences of length 4 (*top*), 5 (*middle*), and 6 (*bottom*), where high error (s.e.m.>20%) history sequences, which correspond to infrequent sequences, have been excluded. Exclusion of these sequences is necessary since, if a sequence is expressed only once it will trivially deterministically evoke a single behavior. Similarly, at low trial numbers, stochasticity is difficult to establish. Insets: Confusion matrices for nonparametric policies of each history length for right and left port choice (*left*) and repeat and switch (*right*). Data is from the 80-20 condition.

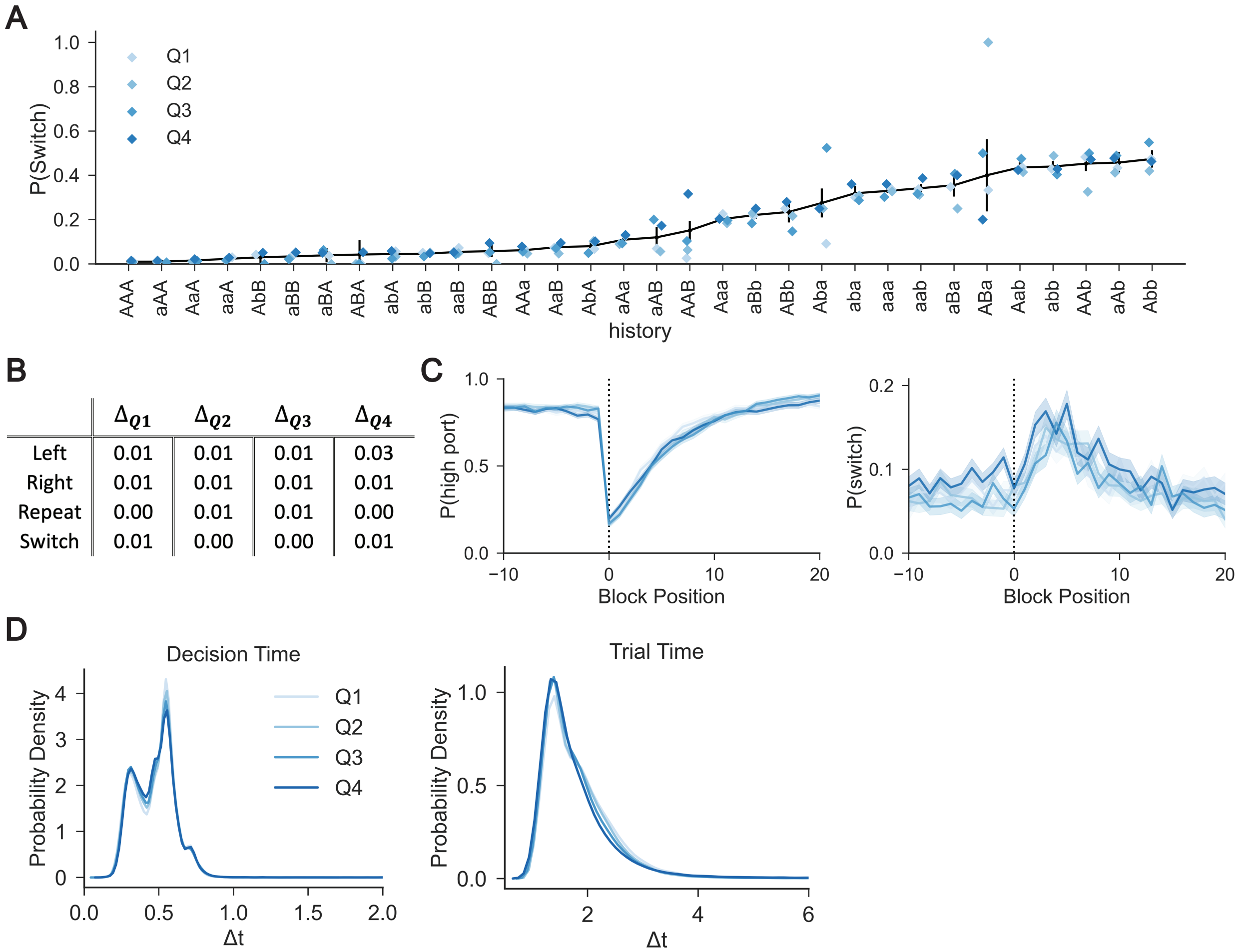


| | $\Delta_{Q1}$ | $\Delta_{Q2}$ | $\Delta_{Q3}$ | $\Delta_{Q4}$ |
|---|---|---|---|---|
| Left | 0.01 | 0.01 | 0.01 | 0.03 |
| Right | 0.01 | 0.01 | 0.01 | 0.01 |
| Repeat | 0.00 | 0.01 | 0.01 | 0.00 |
| Switch | 0.01 | 0.00 | 0.00 | 0.01 |

**Figure 4:** Stationarity of behavioral characteristics within sessions. (A) Conditional switch probabilities for each quartile of a session (Q1 to Q4), overlaid on aggregate switch probabilities (black line). Binomial standard error shown for aggregate conditional switch probabilities. (B) Absolute value differences (Δ) for each action from the confusion matrices in Figure 2E (aggregate data) for each within-session quartile. (C) $P_{high-choice}$ (*left*) and $P_{switch}$ (*right*) as a function of trial number surrounding state transition (block position 0) for each quartile of a session. Dark lines show the mean across trials at the same block position and the shading shows the standard error. (D) Probability density distributions of trial times (in seconds) for each quartile in a session. *Left*: Distribution of time from center port to side port (decision time), *right*: distribution of time from center port to center port (trial initiation to next trial initiation). Probability densities integrate to 1 across the continuous distribution of durations. Data is from the 80-20 condition.

A

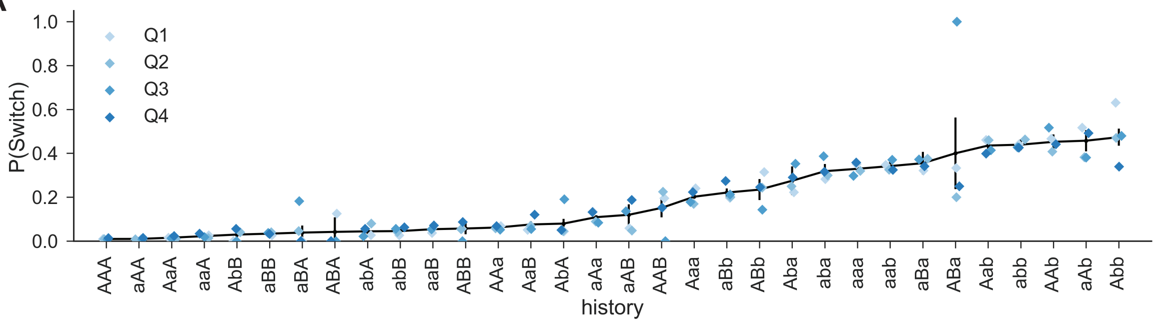


B

| | $\Delta_{Q1}$ | $\Delta_{Q2}$ | $\Delta_{Q3}$ | $\Delta_{Q4}$ |
|---|---|---|---|---|
| Left | 0.00 | 0.01 | 0.00 | 0.02 |
| Right | 0.01 | 0.00 | 0.02 | 0.01 |
| Repeat | 0.01 | 0.01 | 0.01 | 0.00 |
| Switch | 0.03 | 0.00 | 0.00 | 0.01 |

C

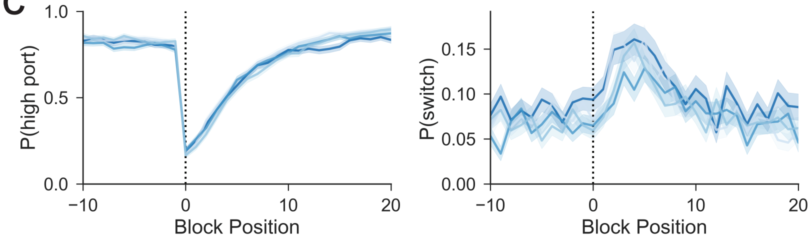


D

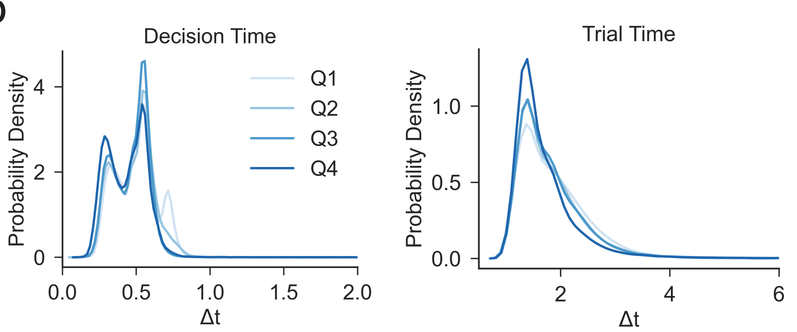


**Figure 5:** Stationarity of behavioral characteristics across sessions. (A) Conditional switch probabilities for session quartiles over the duration of training (i.e., Q1 for early training sessions, Q4 for late training sessions), overlaid on aggregate switch probabilities (black line). Binomial standard error shown for aggregate conditional switch probabilities. (B) Absolute value differences ($\Delta$) for each action from the confusion matrices in Figure 2E (aggregate data) for each across-session quartile. (C) $P_{high-choice}$ (*left*) and $P_{switch}$ (*right*) as a function of trial number surrounding state transition (block position 0) for each across-session quartile. Dark lines show the mean across trials at the same block position and the shading shows the standard error. (D) Probability density distributions of trial times (in seconds) for each across-session quartile. *Left*: Distribution of time from center port to side port (decision time), *right*: distribution of time from center port to center port (trial initiation to next trial initiation). Probability densities integrate to 1 across the continuous distribution of durations. Data is from the 80-20 condition.

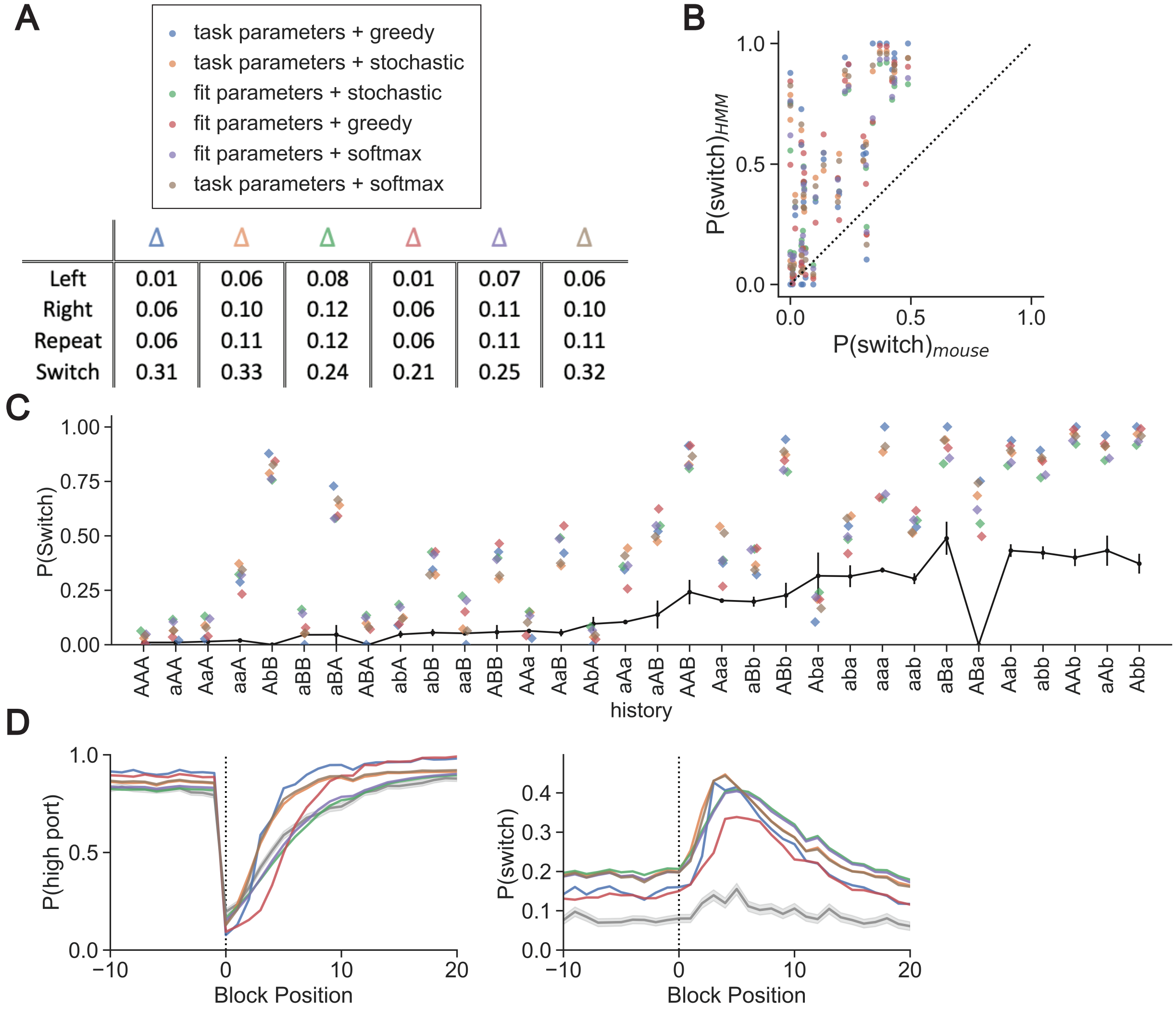


| | Δ | Δ | Δ | Δ | Δ | Δ |
|---|---|---|---|---|---|---|
| Left | 0.01 | 0.06 | 0.08 | 0.01 | 0.07 | 0.06 |
| Right | 0.06 | 0.10 | 0.12 | 0.06 | 0.11 | 0.10 |
| Repeat | 0.06 | 0.11 | 0.12 | 0.06 | 0.11 | 0.11 |
| Switch | 0.31 | 0.33 | 0.24 | 0.21 | 0.25 | 0.32 |

**Figure 6:** Supplementary Figure 6. Alternative parameterizations of Hidden Markov models fail to capture mouse behavior. (A) Absolute values of the differences between each of the HMM's confusion matrices and nonparametric confusion matrix (Fig. 2E). Models, as designated by the labeled indicator colors: an HMM using the true emission and transition probabilities from the task coupled with a greedy policy, stochastic (Thompson sampling) policy, or a softmax policy with intermediate temperature value, as well as an HMM using mouse-fit emission and transition probabilities with each policy. (B) Conditional switch probabilities predicted by each HMM plotted against those observed in mice. Dashed line indicates the unity line. Model indicator colors correspond to those of (A). (C) Conditional switch probabilities predicted by each HMM, overlaid on the conditional switch probabilities of the mouse (black line). Each data point represents the predicted conditional switch probability by the associated HMM for a given history sequence. Error bars show the binomial standard error for the mouse conditional switch probabilities. Histories are sorted as in Fig. 2D. (D) $P_{high-choice}$ (*left*) and $P_{switch}$ (*right*) as a function of trial number surrounding state transition (block position 0) for each HMM (colors, corresponding to (A)), as well as for the observed mouse behavior (gray). Dark lines show the mean across trials at the same block position, and the shading shows the standard error.

**Table 1: Summary statistics for behavioral performance in each of the reward probability conditions.**

| $p$ | P(left) | P(switch) | Rewards/session (mean ± SD) | Trials/session (mean ± SD) | ITI (s) (mean ± SD) | ITI (s) (median ± MAD) | P(lose-switch) | P(win-switch) | Decision Time (s) (mean ± SD) |
|---|---|---|---|---|---|---|---|---|---|
| .90 | 0.49 | 0.083 | 500 ± 116 | 649 ± 138 | 2.21 ± 4.27 | 1.66 ± 1.02 | 0.28 | 0.027 | 0.46 ± 0.14 |
| .80 | 0.51 | 0.067 | 514 ± 77 | 748 ± 104 | 2.05 ± 3.14 | 1.65 ± 0.79 | 0.18 | 0.015 | 0.47 ± 0.14 |
| .70 | 0.47 | 0.067 | 461 ± 71 | 784 ± 110 | 1.89 ± 2.45 | 1.61 ± 0.59 | 0.14 | 0.015 | 0.45 ± 0.13 |

**Table 2: Individual mouse summary statistics for task performance.**

| | 90-10 | | | 80-20 | | | 70-30 | | |
|---|---|---|---|---|---|---|---|---|---|
| Mouse | $P_{high-choice}$ (mean ± SD) | $P_{reward}$ (mean ± SD) | $\tau$ (trials) | $P_{high-choice}$ (mean ± SD) | $P_{reward}$ (mean ± SD) | $\tau$ (trials) | $P_{high-choice}$ (mean ± SD) | $P_{reward}$ (mean ± SD) | $\tau$ (trials) |
| A | 0.86 ± 0.048 | 0.79 ± 0.039 | 3.09 | 0.84 ± 0.039 | 0.70 ± 0.026 | 4.34 | 0.77 ± 0.042 | 0.61 ± 0.028 | 6.09 |
| B | 0.84 ± 0.046 | 0.77 ± 0.038 | 2.87 | 0.83 ± 0.050 | 0.70 ± 0.032 | 3.61 | 0.74 ± 0.052 | 0.60 ± 0.028 | 5.34 |
| C | 0.84 ± 0.041 | 0.77 ± 0.038 | 3.05 | 0.81 ± 0.040 | 0.69 ± 0.024 | 4.69 | 0.73 ± 0.051 | 0.59 ± 0.028 | 5.23 |
| D | 0.86 ± 0.03 | 0.77 ± 0.027 | 3.19 | 0.84 ± 0.049 | 0.70 ± 0.032 | 4.50 | 0.74 ± 0.052 | 0.59 ± 0.022 | 6.26 |
| E | 0.86 ± 0.03 | 0.78 ± 0.030 | 3.14 | 0.83 ± 0.044 | 0.69 ± 0.025 | 4.81 | 0.72 ± 0.060 | 0.59 ± 0.029 | 6.54 |
| F | 0.85 ± 0.04 | 0.78 ± 0.038 | 2.49 | 0.82 ± 0.050 | 0.69 ± 0.030 | 3.71 | 0.68 ± 0.063 | 0.58 ± 0.025 | 6.82 |

**Table 3: Log-likelihoods of held-out data given each model. The nonparametric model uses history length 3, and the logistic regression is the reduced logistic regression with six input features. 'HMM, true' refers to the Thompson sampling HMM with the true task parameters, and 'HMM, fit' refers to that with the mouse-fit parameters.**

| Models | Held-out log-likelihood |
|---|---|
| Nonparametric | -0.180 |
| RFLR | -0.182 |
| F-Q Model | -0.182 |
| Sticky HMM | -0.182 |
| Logistic regression | -0.182 |
| HMM, fit | -0.325 |
| HMM, true | -0.359 |
